# Single-cell immune profiling reveals novel thymus-seeding populations, T cell commitment, and multi-lineage development in the human thymus

**DOI:** 10.1101/2022.02.18.481026

**Authors:** Martijn Cordes, Kirsten Canté-Barrett, Erik B. van den Akker, Federico A. Moretti, Szymon M. Kiełbasa, Sandra Vloemans, Laura Garcia-Perez, Cristina Teodosio, Jacques J.M. van Dongen, Karin Pike-Overzet, Marcel J.T. Reinders, Frank J.T. Staal

## Abstract

T cell development in the mouse thymus has been studied rather extensively; in contrast, strikingly little is known regarding T cell development in the human thymus. To close this knowledge gap, we used a combination of single-cell techniques and functional assays to perform deep immune profiling of human T cell development, focusing on the initial stages of pre-lineage commitment. We identified three thymus-seeding progenitor populations that also have counterparts in the bone marrow. In addition, we found that the human thymus physiologically supports the development of monocytes, dendritic cells, and NK cells, as well as limited development of B cells. These results are an important step towards monitoring and guiding regenerative therapies in patients following hematopoietic stem cell transplantation.

## Introduction

T cell development occurs in the thymus, while all other blood cell lineages develop primarily in the bone marrow (BM) and/or spleen. The thymus consists of a mixture of thymic stromal cells and developing thymocytes, and crosstalk between these cell types provides the appropriate developmental signals needed for mature T cells to develop. T cell development follows a series of distinct phenotypic stages characterized by the expression of specific surface receptors, notably CD4 and CD8. In both humans and mice, thymocyte development proceeds through successive stages, starting with CD4^-^CD8^-^ double-negative (DN) cells, followed by CD4^+^CD8^+^ double-positive (DP) cells, and ending with CD4^+^CD8^-^CD3^+^ or CD8^+^CD4^-^CD3^+^ single-positive (SP) cells. The DN subset can be further subdivided into four consecutive stages in mice (DN1 through DN4) and three consecutive stages in humans (DN1 through DN3). In addition to differences in surface marker expression, mouse and human DN subpopulations also differ with respect to the rearrangement status of their T cell receptor (TCR) genes and their dependence on cytokines ^1, 2^. In humans, the three DN stages are traditionally identified based on their expression profile consisting of the stem cell markers CD34, CD38, and CD1a (Weerkamp et al., 2006). CD34 is a marker for hematopoietic stem cells and progenitors of all hematopoietic lineages; in the thymus, the most immature cells express CD34, and this expression then declines as the cells mature. Concomitant with this decrease in CD34 expression, thymocytes first express CD38, followed by CD1a, roughly coinciding with full αβ T cell commitment ^3^. Thus, the developing human DN compartment is typically subdivided into the CD34^+^CD38^-^CD1a^-^ (DN1) stage (representing the most immature thymocyte subset) followed by the CD34^+^CD38^+^CD1a^-^ (DN2) and CD34^+^CD38^+^CD1a^+^ (DN3) stages.

Using DNA microarrays, we previously found that the above-mentioned stages in human T cell development coincide with precisely regulated patterns of TCR gene rearrangement and upregulation of T cell lineage specific genes ^1, 2^. In addition, our group and others showed that the CD34^+^CD1a^-^ subsets contain progenitors with B lymphoid, myeloid, dendritic, NK cell, and even erythroid lineage potential ^4, 5, 6, 7^; however, whether any of these cell lineages actually develops within the healthy human thymus is currently unknown.

Another open question—even in mice—is the precise nature of the cells that enter the thymus ^8^. Hematopoietic progenitors from the BM enter the circulation and migrate to the thymus, where they commit to the T cell lineage. Thymic progenitors are continuously outcompeted by newly imported progenitors from the BM, an essential process that maintains healthy T cell development ^9^. However, the precise identity of these so-called thymus-seeding progenitor (TSP) cells is controversial, as some BM progenitor subpopulations may already possess T lineage potential ^10, 11^; moreover, their extremely low numbers hamper their identification. Here, we focused on the most immature DN1 subset, which presumably contains the non-committed cells that recently seeded the thymus and may already have early specification for the T cell lineage.

Although genetic loss-of-function (LOF) studies in mice have been used to dissect thymocyte developmental trajectories and checkpoints, similar studies in humans are sparse. The closest study of genetic LOF in human T cell development used cells that were obtained from various patients with severe combined immunodeficiency (SCID) and xenotransplanted into NSG (NOD scid gamma) mice, revealing increased heterogeneity and—unexpectedly—novel checkpoints in human T cell development ^12^.

Single-cell RNA sequencing (scRNA-seq) can be used to identify and characterize rare thymocyte populations and visualize the transcriptional landscape at the single-cell level, providing new insights into T cell development trajectories. Recently, several studies used scRNA-seq to show that most developmental stages based on surface markers are not linear and homogeneous, but rather have overlapping transcriptional programs, resulting in a heterogeneous continuum ^13, 14, 15^.

To examine cellular heterogeneity in the human thymus, we performed scRNA-seq and corresponding V(D)J-enriched scRNA-seq in several sorted subsets with overlapping developmental potential. We then performed spectral flow cytometry of developing human thymocytes in order to perform a detailed phenotypic analysis. Specifically, we characterized: *i*) the nature and differentiation of TSP subpopulations, *ii*) T cell lineage commitment, and *iii*) the development of non-T cell lineages in the human thymus. To complement our scRNA-seq analyses, we also used functional assays and lineage tracing and identified three distinct TSP subpopulations, several non-T cell developmental pathways in the human thymus, and branching points between αβ and γδ T cells, including a stepwise commitment towards the αβ T cell fate.

## Results

### Generation of a single-cell atlas of human T cell development

To profile the complete thymocyte population, including its rare subsets, we performed scRNA-seq on sorted thymocytes obtained from six healthy donors (**Fig**. 1A), focusing on early CD4^-^CD8^-^ double-negative (DN) thymocytes. We focused specifically on this early stage in order to gain insight into the nature of the progenitor cells that seed the thymus, the T cell specification and commitment processes, and possible non-T cell lineage differentiation. We therefore used flow cytometry to sort DN1 (CD34^+^CD38^-^CD1a^-^), DN2 (CD34^+^CD38^+^CD1a^-^), and DN3 (CD34^-^CD38^+^CD1a^+^) thymocytes (Dik et al., 2005) (**Fig**. 1B and **Suppl. Fig**. S1A). We also analyzed several additional sorted subsets, including CD4^+^ immature single-positive (ISP), CD4^+^CD8^+^ double-positive (DP), CD4^+^ SP, and CD8^+^ SP thymocytes. The DP subset was further subdivided into CD4^+^CD8^+^CD3^-^ and CD4^+^CD8^+^CD3^+^ thymocytes (**Fig**. 1B and **Suppl. Fig**. S1B). From these 8 sorted populations we generated 8 scRNA-seq libraries for gene expression profiling using the 10x Genomics platform. In addition, these 8 sequencing libraries were used for targeted amplification of both the αβ and γδ V(D)J genes using TCR-specific primers, resulting in 16 additional libraries for TCR sequencing ^16^. In total, we obtained a dataset for approximately 40,000 cells obtained from six donors (**Fig**. 1 C, D), with both gene expression data and αβ and γδ TCR rearrangement data for each cell.

**Fig. 1.**
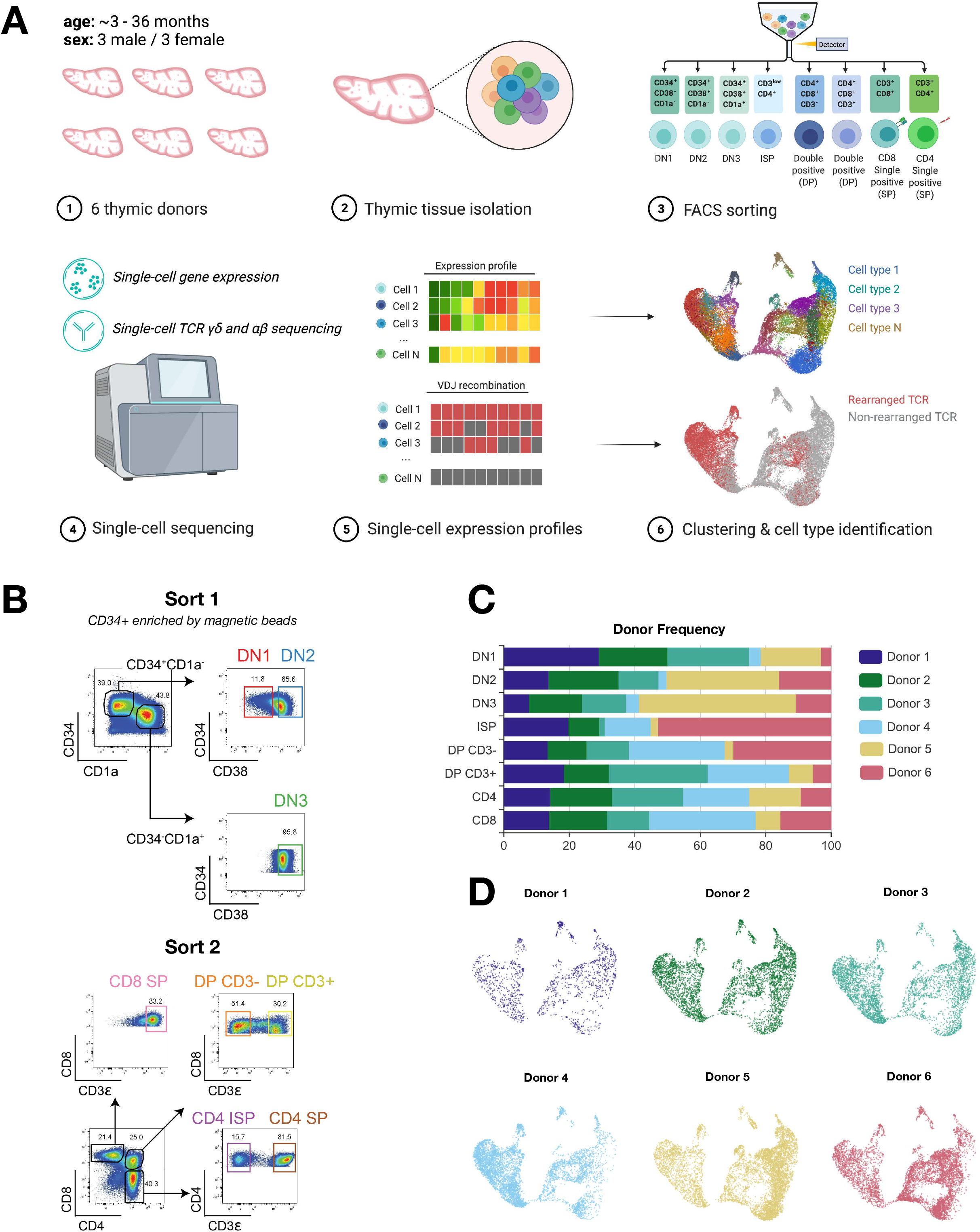
Single-Cell Transcriptomics of Human Thymocytes. (A) _+_Experimental scheme: thymocytes from 6 different donors were isolated, pooled and sorted for 8 sub-populations representing 8 differentiation stages of human T cell development. All 8 subpopulations were sequenced for gene-expression profiling and TCR analysis. DN: double-negative, ISP: CD4 immature single-positive, DP: double-positive. (B) Sorting strategies of the 3 DN populations after CD34_+_ enrichment (sort 1), and the 5 DP/SP populations (sort 2). (C) Donor proportion per developmental stage. (D) Contributions of each individual donor to total UMAP projection (Fig 2).

### Overall analysis places our sorted fractions at the expected developmental stages and reveals various stages in which γδ T cells and plasmacytoid dendritic cells continue to develop

After *in silico* quality control to remove low-quality cells and doublets ^17^, we visualized the overall scRNA-seq data using a two-dimensional UMAP (uniform manifold approximation and projection) to capture global transcriptional changes in developing thymocytes. We found that our high-quality transcriptome data clustered as a continuum spanning the sorted subsets (**Fig**. 2A). Moreover, gene expression profiles were consistent with known changes that occur during human T cell development (**Fig**. 2B), with CD34 expression reflecting the most immature thymocytes that appear shortly after entering the thymus. Notch signaling subsequently upregulates *CD7* expression, marking specification towards the T cell lineage. The first T cell specific target of the Notch signaling pathway, TCF1 (encoded by the *TCF7* gene), is then expressed, followed by *BCL11B*, both of which are critical transcription factors involved in committing early thymocytes to the T cell lineage in mice ^18, 19, 20^. A subset of early thymocytes with high *BCL11B* expression also express *CD1A* and *PTCRA* (**Fig**. 2B), marking T cell commitment and the initiation of TCR rearrangement. After TCRβ selection in DP cells, the cells differentiate further into SP cells. Interestingly, we noticed that after correcting for cell cycle, the DN cells were either in a state of proliferation (expressing *TOP2A*) or recombination (expressing *RAG1*), confirming that these two processes occur at distinct times during development (**Fig**. 2C). The timing of TCR rearrangement can be illustrated by aligning the γδ (**Fig**. 2D) and αβ TCR recombination data (**Fig**. 2E) to the UMAP plot, showing that γδ rearrangements occur as early as the DN2 stage and through the ISP and even DP stages. The ISP stage that marks the peak of TCRβ rearrangements coincides with the proliferation of *TCRB*-rearranged cells, while the majority of *TCRG*-expressing cells follow the path with ongoing recombination (**Fig**. 2D, E).

**Fig. 2.**
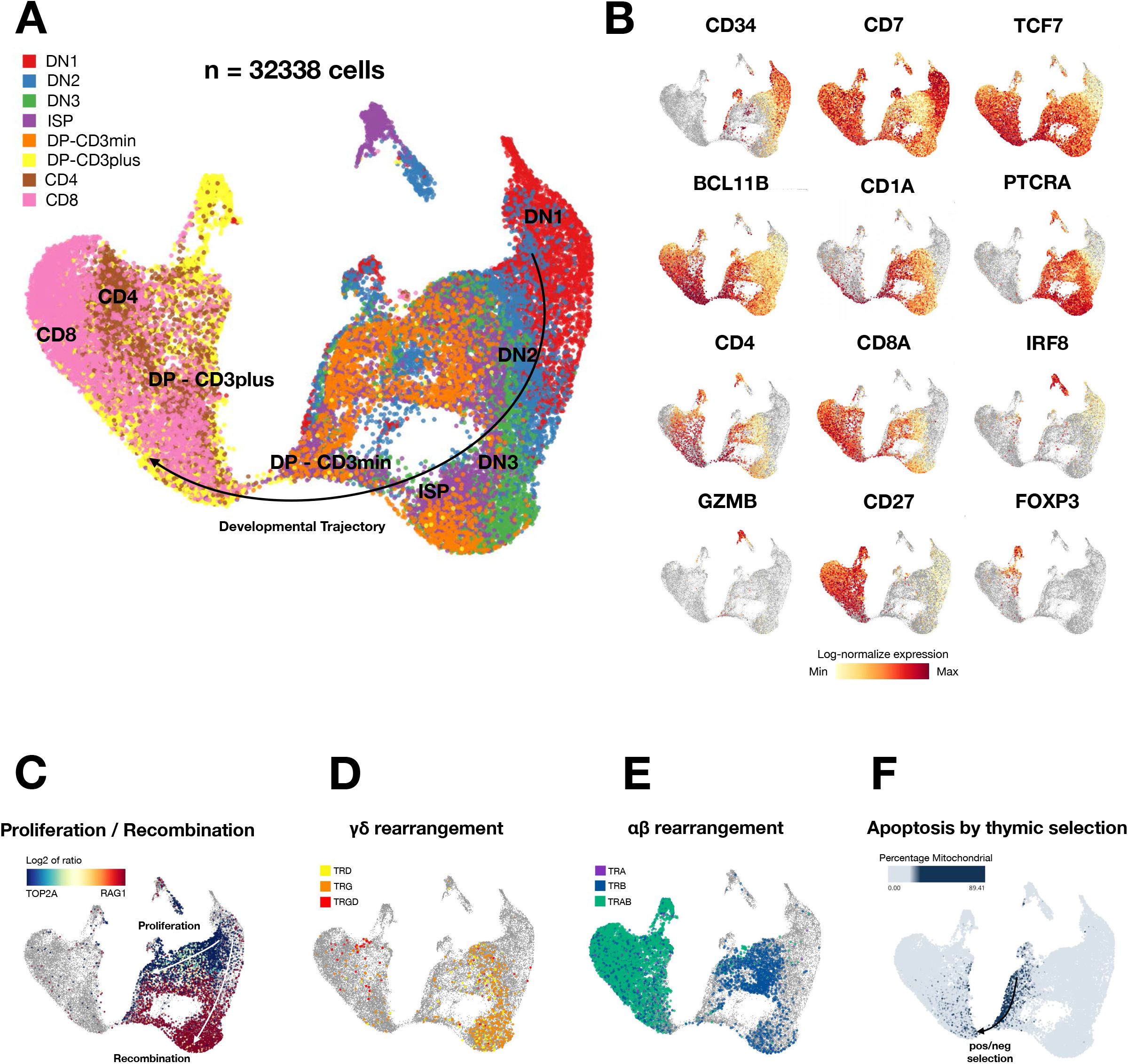
UMAP projections of all 8 sorted populations reveals gene expression related to cellular differentiation and biological processes. (A) UMAP projections of complete scRNA-seq datasets colored by sorted populations that reflect T cell-developmental stages. (B) Gene expression level (red is high expression) of key surface markers and transcription factors projected on the UMAP. (C) *TOP2A* (blue) and *RAG1* (red) gene expression indicating proliferation and recombination, respectively. (D) Cells with rearranged TCRγ (orange), TCRδ (yellow), or fully rearranged TCRγδ (red). (E) Cells with rearranged TCRα (purple), TCRβ (blue), or fully rearranged TCRαβ (green). (F) Mitochondrial expression (blue) concomitant with cells in apoptosis during thymocyte selection predominantly just prior to the major transition between DP, CD3 to DP, CD3 _+_ stages.

Recently, several studies using scRNA-seq reported intrathymic development of plasmacytoid dendritic cells (pDCs) ^21, 22^. Consistent with these findings, we also detected cells with a pDC expression profile—i.e., expressing *IRF8* (**Fig**. 2B), *TCF4*, and *SPIB* (**Suppl. Fig**. S2)—within both the DN2 and ISP thymic subpopulations; these cells clustered above the main UMAP plot (**Fig**. 2B and **Suppl. Fig**. S2). Importantly, the ISP-derived pDCs could be distinguished from the DN2-derived pDCs by their expression of *PTCRA, CD4*, and *GZMB* (**Fig**. 2B), suggesting that pDCs either arise from distinct progenitor cells or co-develop with T cells ^23^.

Apoptosis occurs in developing thymocytes after specific checkpoints, and cells undergoing apoptosis can be identified by measuring the percentage of reads mapping to the mitochondrial genome per cell (**Fig**. 2F) ^17, 24^. As expected, *TCRB*-rearranged cells that have undergone TCRβ selection followed by rearrangement of the *TCRA* gene (**Fig**. 2E) subsequently undergo either positive or negative selection ^25^ during the time of *CD3* expression in DP cells (Dik et al., 2005). Negative selection of autoreactive thymocytes accounts for the loss of >90% of thymocytes via apoptosis ^26^. After crossing this tight bottleneck—indicated by the narrow “bridge” connecting the right and left sides of the UMAP plot—positively selected thymocytes continue their development to the CD3^+^ DP stage. Cells expressing *CD27* can be rescued from negative selection and become regulatory T cells (Tregs), indicated by expression of the transcription factor *FOXP3* ^27^; this is also visible in our dataset (**Fig**. 2B and **Suppl. Fig**. S2).

### Identification of three thymus-seeding progenitor cell populations

Hematopoietic stem cells (HSCs) that migrate from the bone marrow to the thymus initiate the differentiation process shortly after entering the thymic environment. Thus, naïve TSPs that still resemble HSCs are extremely rare. Moreover, whether partially differentiated lymphoid progenitors from the circulation can initiate T cell development in the thymus is currently unclear ^28^. To identify these early progenitors within the context of early thymic development, we re-analyzed the ∼10,000 DN1, DN2, and DN3 cells and generated a new UMAP plot (**Fig**. 3A and **Suppl. Fig**. S3A). We found that the DN1, DN2, and DN3 cells generally followed their expected developmental order (**Fig**. 3B, C and **Suppl. Fig**. S3B); moreover, cluster marker analysis confirms heterogeneity among the DN cell populations (**Fig**. 3D). The cells in clusters 1-5 have an expression profile that suggests strong multi-lineage potential based on their expression of the multipotency markers *HOPX* and *ACY3*, the B cell lineage related genes *IGHM, MEF2C*, and *IGLL1*, and the myeloid marker *MPO* (**Fig**. 3D). The multi-lineage potential of clusters 1 and 2 was also confirmed by their lack of expression of the key Notch signaling induced transcription factors *TCF7* and *BCL11B* _19_

**Fig. 3.**
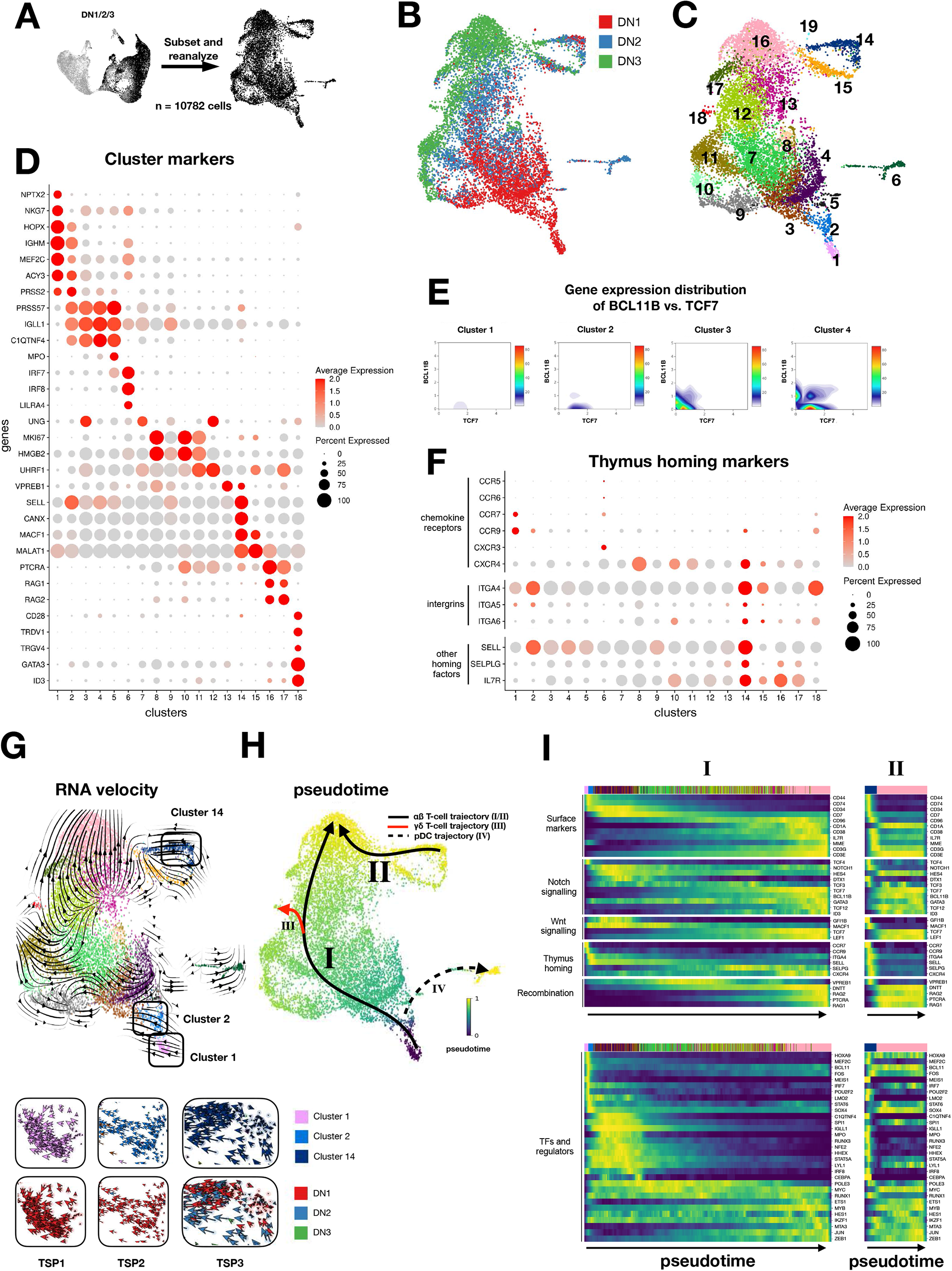
Reanalysis of CD34+ progenitor populations reveals multiple developmental trajectories. (A) Reclustering of DN1, DN2, and DN3 cells. *(A) Reclustering of DN1, DN2, and DN3 cells* (B) UMAP projection of DN1 (red), DN2 (blue), and DN3 (green) cells. (C) UMAP projection of 18 clusters identified by cluster analysis. (D) Dot plot depicting the genes that identify clusters based on differential gene expression. (E) Density plots of TCF7 and BCL11B gene expression in clusters 1-4. (F) Dotplot indicating relative gene expression of thymus-homing markers per cluster. (G) Prediction of the global differentiation of thymocytes by RNA velocity analysis (top). TSP1,2, and 3 have been enlarged and indicated for DN1, 2, or 3 origin (bottom). Arrows indicate differential splicing activity between TSPs. Global velocity streamlines are visualized by the large arrows which is a summary of all single cell velocities (small arrows, bottom inlays). (H) Pseudo-temporal ordering of cells colored from dark purple to yellow (0-1) according to maturation state. Lines represent differentiation trajectories of two routes of αβ T cell development (black lines), γδ T cell (red line) and pDC (dotted line) development. (i) Heatmap showing gene expression of surface markers, signaling pathways, thymus homing markers, recombination genes and transcription factors involved in the T cell differentiation ordered according to the pseudo-time of the conventional (I) and alternative (II) αβ T cell development trajectories displayed in (H).

The earliest DN cells (i.e., in cluster 1) lack *CD7* expression, indicating that the Notch signaling pathway is not yet activated in this stage. Indeed, the cells in cluster 2 are the first to show *CD7* expression, resulting in low expression of *TCF7*, followed by an upregulation of both *TCF7* and *BLC11B* in clusters 3 and 4 (**Fig**. 3E). We then searched for rare, novel thymocyte subpopulations—including TSPs—in our DN dataset. TSPs enter the lobes of the thymus via high endothelial venules ^29, 30^, then home directly to the thymic environment by interacting with ligands expressed by thymic epithelial cells ^31^. We therefore searched our dataset for cell populations that express chemokine receptor genes (*CCR7, CCR9, CXCR3*, and *CXCR4*) ^32, 33, 34, 35^, integrins (*ITGA4, ITGA5*, and *ITGA6*), and other homing factors *(CD62L/SELL, CD162L/SELPG*, and *CD127/IL7R)*^28, 31^. Based on these gene sets, the cells assigned to clusters 1, 2, and 14 seem to have the most thymus-homing activity; we named these thymus-seeding populations TSP1, TSP2, and TSP3, respectively (**Fig**. 3F, G). Notably, the cells in TSP1-3⍰ were derived primarily from the most immature sorted population, namely the DN1 subpopulation (**Fig**. 3B, G); therefore, our downstream analyses of the three TSP subsets were performed using data obtained from DN1 thymocytes.

### CD34^+^ progenitor cells follow several developmental trajectories, including an alternative trajectory from progenitors to T cell lineagelllcommitted thymocytes

T cell precursors progress through a series of distinct developmental stages marked by gradual phases of lineage specification, followed by a T cell specific transcriptional program. Before becoming fully committed to the T cell lineage, T cell precursors gradually lose their non-T cell lineage potential ^18, 36^. To gain new insights into the transcriptional changes that accompany lineage commitment in early progenitor populations, we examined the complete DN dataset. The cells in cluster 6 were classified as pDCs based on their expression of *IRF8* and *LILRA4* (also known as *CD85g*). Based on their expression of the proliferation markers *MKI67* and *HMGB2*, clusters 7-12 were classified as proliferating cells. Interestingly, we found that cluster 13 had high expression of the pre-B cell receptor gene *VPREB1*, while cluster 14 (i.e., the TSP3 subset) had high expression of the Wnt pathway associated gene *MACF1* and the homing marker *SELL* (also known as *CD62L*). The cells in cluster 16 were classified as T lineage committed thymocytes based on their high expression of the recombination-activating genes *RAG1* and *RAG2*, as well as expression of *PTCRA* (**Fig**. 3D, **Suppl. Fig**. S3B) and the presence of *TCRB*-rearranged cells (**Suppl. Fig**. S3C). Lastly, we identified a cluster of mature γδ T cells based on the expression of the γδ-associated transcription factors *ID3* and *GATA3* (**Fig**. 3D), as well as expression of *TRDV* and *TRGV* combined with the presence of *TCRG*- and *TCRD*-rearranged cells (**Suppl. Fig**. S3D).

Application of RNA velocity ^37, 38^ and diffusion pseudotime (DPT) ^39^ provided valuable insights into the trajectory of developing thymocytes. We found that overlaying the RNA velocity vectors on the UMAP plot (**Fig**. 3G) predicted the global differentiation of thymocytes into αβ T cells (cluster 16), γδ T cells (cluster 18), or pDCs (cluster 6). Moreover, ordering of the cells by DPT (**Fig**. 3H) relative to cluster 1 confirmed that cells in cluster 6 (i.e., pDCs) and cluster 16 (i.e., committed thymocytes) were the most mature cells in our dataset. Together, these analyses suggest the presence of four major differentiation trajectories in our dataset. The first trajectory is the conventional αβ T cell development route (indicated as route I in **Fig**. 3H) from the most immature, multipotent clusters 1 and 2 (TSP1 and TSP2 cells, respectively) to the rearranged, committed thymocytes in cluster 16. Surprisingly, however, we identified an alternative T cell trajectory (indicated as route II in **Fig**. 3H) that also ends at cluster 16 but starts from the relatively more mature cluster 14 (TSP3 cells). A third trajectory leading to γδ T cells splits off from the first trajectory (route III indicated by the red line, exiting route I in **Fig**. 3H) and ends in cluster 18. Finally, as discussed above, we detected two distinct pDC populations (see **Fig**. 2A, B), one of which is located in the DN2 population; thus, the fourth trajectory revealed by our DPT analysis is used by cells that differentiate into pDCs (route IV indicated by the dashed line in **Fig**. 3H).

### Transcriptional regulation during the early conventional (route I) and alternative (route II) trajectories of αβ T cell development

To gain additional insight into the transcriptional changes that occur during the conventional and alternative trajectories of αβ T cell development, we ordered the DN1, DN2, and DN3 cells along these two routes from the early progenitors in clusters 1 and 2 to committed thymocytes in cluster 16 (i.e., route I) and from the cells in cluster 14 to committed thymocytes in cluster 16 (i.e., route II), as predicted by the RNA velocity and DPT analysis (**Fig**. 3G, H). We also identified differentially activated transcription factors and regulons (transcriptionally co-regulated operons) in each cluster using single-cell regulatory network inference and clustering (SCENIC) ^40^, revealing clear changes in regulatory activity in the T cell differentiation trajectories (**Suppl. Fig**. S3E). For example, the cells in cluster 1 express stem cell like and progenitor-like genes such as the hematopoietic regulatory genes *HOXA9, MEF2C, BCL11A, MEIS1, HOPX, and NPTX2*, as well as the chemokine receptors CCR7 and CCR9, which are important for homing to the thymus. The stemness genes are the first genes to be downregulated following Notch expression (**Fig**. 3D, I and **Suppl. Fig**. S3E). In addition to Notch signaling, Wnt signaling is also active in the DN stages, indicated by the upregulation of *LEF1* even before *TCF7* expression ^41^ (**Fig**. 3I, **Suppl. Fig**. S3F). Interestingly, in these stages *GFI1B* (growth factor independent 1B transcriptional repressor) is also expressed; this protein regulates Wnt/β-catenin signaling, thus indicating TCF/LEF-mediated transcription ^42^. Expression of the master hematopoietic regulator gene *SPI1* (encoding the transcription factor PU.1) remains upregulated through cluster 6, until its expression is finally overridden by the effects of Notch signaling in cluster 7 (**Fig**. 3I and **Suppl. Fig**. S3F). During this early window, multi-lineage transcription factors and regulators are expressed at high levels, including the myeloid-specific genes *MPO, CEPBA*, and *LYZ*, the B cell markers *IGLL1, VPREB3, PAX5*, and *EBF1*, the dendritic markers *IRF8* and *SPIB*, and the erythroid transcription factor *NFE2* (**Fig**. 3I and **Suppl. Fig**. S3F, G). Cells that undergo Notch signaling increase their *HES4* and *DTX1* expression, followed by the expression of *GATA3, TCF7*, and *BCL11B* and a rapid downregulation of *MME* (also known as *CD10*) and *TCF4*. Although the expression of *DNTT* (encoding the enzyme terminal deoxynucleotidyl transferase) starts in cluster 3, its expression increases significantly in cluster 7, coinciding with the increased expression of *TCF7* and *BCL11B* (**Fig**. 3I). At this point, *PTCRA* and CD1A expression marks the cell’s commitment to the T cell lineage. By comparing the predicted transcription factor activity with our clustering and trajectory analyses, we identified novel regulators (**Suppl. Fig**. S3E) with activity during the T cell commitment process (i.e., in clusters 7 through 13). These regulators include: *POLE3*, which is consistent with the recent finding in *Pole3*-deficient mice that T cell differentiation is blocked at the DN3 stage ^43^; *MTA3*, a cell fate regulator involved in B cell differentiation ^44^; and *ZEB1*, which was recently described as playing an essential role in the transition of mouse thymocytes from the DN2 stage to the DP stage ^45^.

Notably, we also found that the pre-B cell receptor gene *VPREB1* is expressed in the earliest cells and continues until the cells begin to express *CD1A* (**Fig**. 3I and **Suppl. Fig**. S3G). In addition, expression of the *RAG* genes together with the expression of *TCF12* (which encodes transcription factor 12, also known as HEB) initiates TCR rearrangement. The cells in cluster 8 are the first to contain a rearranged *TCRG* locus (**Suppl. Fig**. S3D), followed by the completely rearranged *TCRD* locus in clusters 12 and 16, marking the first *TCRB*-rearranged thymocytes (**Suppl. Fig**. S3C).

### Fully rearranged γδ T cells in developing DP thymocytes

We found that *TCRB* rearrangement occurs during the ISP and CD3^-^ DP stages of T cell development, followed by *TCRA* rearrangement in the CD3^+^ DP stage (**Suppl. Fig**. S4A-E). In addition, we found that cells either undergo or have completed their *TCRG* and *TCRD* rearrangement in the ISP and DP stages, including the mature γδ T cells in cluster 16 (**Suppl. Fig**. S4C, F), thus after the initial γδ T cells split off in cluster 18 (**Fig**. 3B, C); indeed CD4^+^CD8^+^ γδ T cells were found in the DP population. This finding suggests that cells can differentiate into γδ T cells at various developmental stages and follow the αβ path for longer than previously anticipated.

### TSP1, TSP2, and TSP3 subsets generally represent HSC-like, MPP-like, and CLP-like thymus-seeding progenitors, respectively

As TSPs originate from the BM, we sought to determine the BM progenitors that give rise to the three TSP subsets that we identified. To this end, we mapped the TSP1, TSP2, and TSP3 cells in the DN1 dataset to a large multimodal dataset that includes all major BM-derived blood cell types and progenitors ^46^ (**Fig**. 4A, B and **Suppl. Fig**. S5). We also sub-annotated the CD34^+^ progenitor population in this dataset using the recently reported BM progenitor markers ^47^. This transcriptomics-based mapping approach revealed general correspondence between the three TSP clusters and distinct BM progenitor populations. Specifically, the majority of TSP1 cells match the HSC annotation, the majority of TSP2 cells match the MPP (multipotent progenitor) annotation, and the majority of TSP3 cells match the LMPP (lympho-myeloid primed progenitor) and CLP (common lymphoid progenitor) annotations (**Fig**. 4C-E); importantly, our TSP3 subset resembles CD34^+^CD7^+^ CLPs previously reported in humans ^48^.

**Fig. 4.**
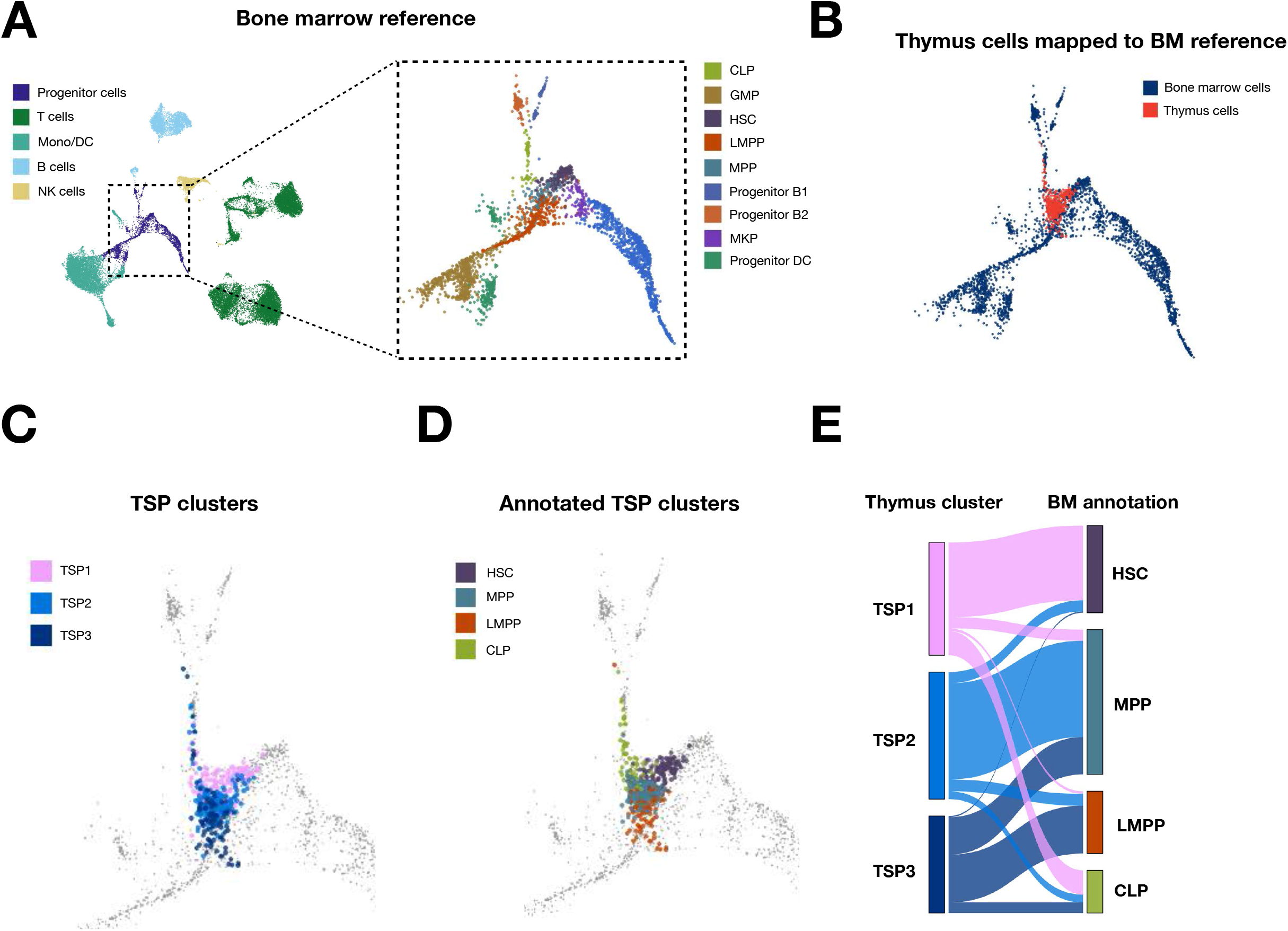
Identification of three thymus-seeding progenitor (TSP) populations. (A) Annotated multimodal BM reference UMAP ^46^ with a focus on progenitor cell populations (outlined by the dotted square). (B) The three thymocyte TSP populations (red) mapped onto the BM progenitor cell (blue) reference UMAP. (C) Thymocyte TSP1 (pink), TSP2 (blue), and TSP3 (dark blue) mapped separately. (D) The same TSPs colored for BM progenitor label after reference mapping. (E) TSPs linked to their BM annotation after reference mapping in (D).

Next, we analyzed differential gene expression between the three TSP subsets and identified subset-specific markers (**Fig**. 5A). TSP1 is a relatively rare CD34^+^CD7^-^ HSC-like population that lacks *CD7* expression but expresses several immature stemness-like genes, including *HOXA9, HOPX, MEF2C*, and *NPTX2* (**Fig**. 5A; see also cluster 1 in **Fig**. 3D). Strikingly, this population seems to be quiescent based on the lack of expression of proliferation genes (**Suppl. Fig**. S3B) and is therefore not a rapidly expanding TSP population. TSP2 is a CD34^+^CD7^+^ MPP-like population that expresses *CD7* but no other T cell specific genes; in contrast, these cells express genes in non-T cell lineages, including *IGHM, IGLL1, NFE2, MPO, IRF8, LY6E, MME*, and *SPI1*, suggesting that they retain their multipotency (**Fig**. 5A; see also cluster 2 in **Fig**. 3D); although expression of these genes continues through clusters 3-5, these clusters also begin to express T cell genes such as *NOTCH1, HES4*, and *LEF1*. Finally, TSP3 is a CD34^+^CD7^+^ CLP-like population that also expresses the homing marker genes *CD62L* and *CXCR4* and Wnt pathway genes such as *MACF1* ^49^ (see also cluster 14 in **Fig**. 3D, F). Given the high expression of *IL7R* and other lymphoid marker genes and the low expression of non-T cell markers, TSP3 cells appear to be primed to develop rapidly into T cells upon entering the thymus.

**Fig. 5.**
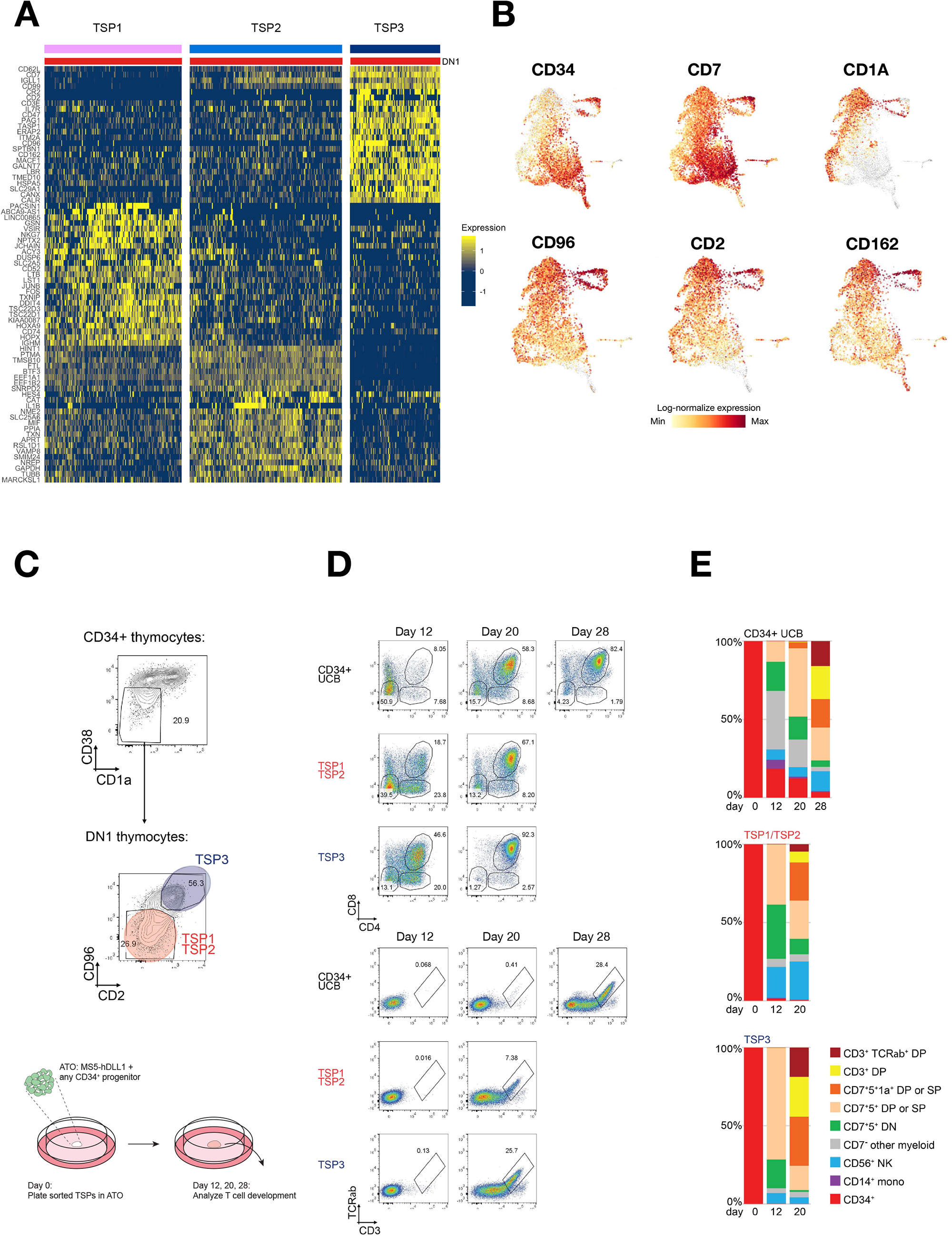
TSP1, 2 and TSP3 differ in T-cell differentiation kinetics *in silico* and in vitro. (A) Heatmap of differential gene expression between the three identified TSPs (with DN1 cells). (B) Examples of gene expression of phenotypic markers of TSPs. (C) Flow sorting strategy of TSPs (within CD34 CD38 CD1a DN1 thymocytes) using CD2 and CD96 as distinguishing markers. Sorted TSPs were cultured in an artificial thymic organoid (ATO). (D) Flow cytometry analysis of TSP cells and CD34 -enriched umbilical cord blood (UCB) cells after 12, 20 and 28 days (only CD34 UCB cells) of culture in an ATO. (E) Bar graphs indicating various cell type proportions of sorted populations after 12, 20, and 28 days of ATO culture.

### TSP1/2 cells and TSP3 cells have distinct T cell differentiation kinetics based on in silico data

Based on our *in silico* DPT analyses (**Fig**. 3H and I), we predicted that the two trajectories to reaching αβ T cells in cluster 16 (i.e., trajectory I from clusters 1 and 2, and trajectory II from cluster 14) follow distinct time courses. When we focused on the RNA velocity data for the various TSP subsets, we found differences in splicing activity (reflected by differences in the length of the arrows) (**Fig**. 3G, insets). Compared to the cells in TSP1, the cells in TSP2 have relatively high splicing activity (i.e., longer arrows); moreover, the arrows in TSP2 have clear directionality. In contrast, the cells in TSP3 have virtually no splicing activity at their putative starting point in the topmost right corner of cluster 14 (**Fig**. 3G, bottom insets), whereas the neighboring cells in cluster 14 have high splicing activity, suggesting the onset of differentiation. We then ordered the cells along the two predicted trajectories (with TSP1 and TSP3 cells as the starting points for trajectories I and II, respectively) and plotted the gene expression data on a pseudotime axis (**Fig**. 3I). We found that the two trajectories follow a similar gene expression pattern but differed considerably with respect to their total duration; specifically, the TSP1 cells first differentiated into several consecutive intermediate cell types (identified as clusters 3 through 8) before becoming T cell lineage committed thymocytes, while the TSP3 cells reached this stage far more rapidly.

### *In vitro* data collected using artificial thymic organoids support our in silico findings regarding the more rapid differentiation of TSP3 cells compared to TSP1/2 cells

To confirm our *in silico* results suggesting that TSP3 cells are primed to develop more rapidly into T cells upon entering the thymus compared to the less mature TSP1 and TSP2 subsets (which take longer to reach cluster 16), we isolated and cultured these subpopulations. Using flow cytometry, we investigated all of the cell surface markers that were differentially expressed between the three TSP subsets (**Fig**. 5A). We found that the CD34^+^CD38^-^CD1a^-^ DN1 thymocytes contained a distinct CD2^-^CD96^-^ subpopulation, the majority of which was CD7^-^ and CD162^+^, consistent with TSP1/TSP2 cells; in contrast, the CD2^+^CD96^+^ subpopulation was CD7^+^, CD162^+^, and IL7R^+^, consistent with TSP3 cells (**Fig**. 5B, C and data not shown). We then tracked the development of flow-sorted CD2^-^CD96^-^ DN1 (i.e., TSP1/TSP2) and CD2^+^CD96^+^ DN1 (i.e., TSP3) cells in artificial thymic organoids ^50^ cultured for up to 28 days (**Fig**. 5C). As a control, we found that when starting with CD34^+^ cells isolated from cord blood, only half of the cells were CD7^+^CD5^+^ by day 12; in contrast, virtually all of the TSP1/2 and TSP3 cells were CD7^+^CD5^+^ by that same time point (**Fig**. 5D, E and **Suppl. Fig**. S6). CD4^+^CD8^+^ DP cells appeared by day 12 but were far more pronounced when starting with TSP3 cells compared to TSP1/2 cells. Finally, approximately 25% of the TSP3 cells were fully committed αβ T cells by day 20, compared to only 7% of TSP1/2 cells and 0% of CD34^+^ control cells (**Fig**. 5D, E). Thus, these *in vitro* data support our prediction that TSP3 cells are indeed primed to develop much more rapidly into T cells compared to TSP1 and TSP2 cells.

### Development of other, non-T cell types in the thymus

To gain more insight into the multi-lineage potential of thymocytes, we integrated our complete thymus dataset with two different human BM datasets; one dataset contains 40,000 mature BM cells from the immune census dataset ^51^, and the other dataset contains 25,000 early hematopoietic progenitor cells from a CD34-enriched BM dataset ^52^ (**Fig**. 6A and **Suppl. Fig**. S7A). All three datasets were processed, integrated, and annotated (for details, see STAR Methods), resulting in a combined dataset containing a total of approximately 77,500 cells. We then generated a three-dimensional UMAP plot using Seurat; the plot was then loaded into the Bioturing Browser ^53^, which makes it possible to rotate the UMAP plot and visualize the cells at different angles (**Fig**. 6 and **Suppl. Fig**. S7). Importantly, this integrated single-cell dataset fully recapitulated the differentiation of HSCs via progenitor populations into a wide range of mature immune cell subsets in the BM (**Fig**. 6B). Notably, the thymocytes that were integrated into the BM setting still followed the overall T cell differentiation trajectory from DN cells through ISP, DP, and finally SP cells (**Fig**. 6C).

**Fig. 6.**
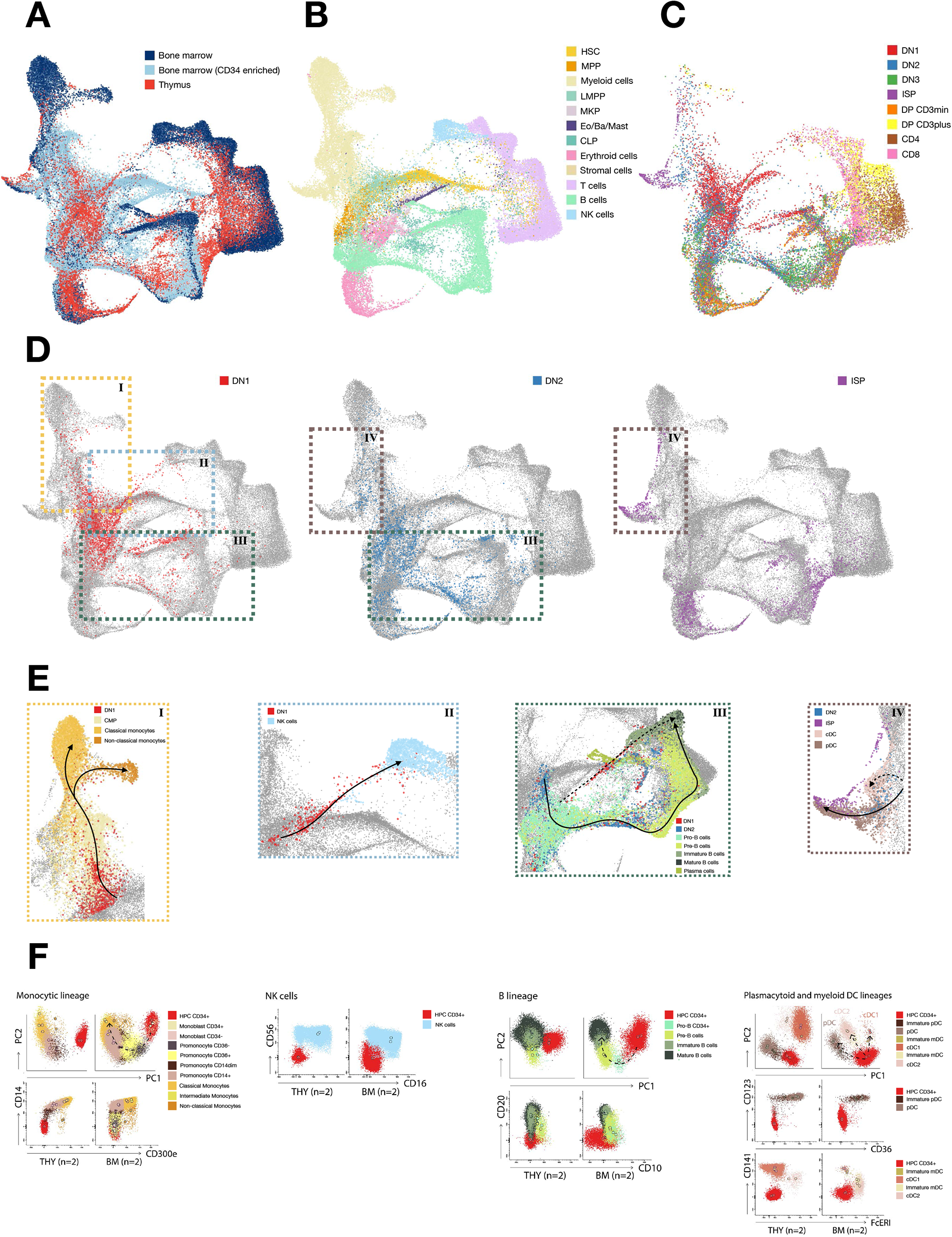
Intrathymic development of alternative lineages. (A) UMAP projection after integration of mature BM cells ^51^ with CD34+ BM ^52^ and our complete Thymus dataset. (B) UMAP projection of annotated BM cells. (C) All eight sorted thymocyte subsets projected on the integrated UMAP. (D) Projection of DN1 (red), DN2 (blue) and ISP cells (purple) on the integrated UMAP. (E) Insets from (D): DN1, DN2 and ISP cell differentiation towards monocytes, natural killer (NK) cells, B cells, classical dendritic cells (cDCs) and plasmacytoid dendritic cells (pDCs). (F) Principal component (PC) analyses and example flow plots (bottom rows) of 28-color spectral flow cytometry of human thymocytes (n=2) and human BM (n=2) to indicate mature non-T cell types (monocytes, NK cells, B cells, cDCs and pDCs) and their progenitors. mDC: myeloid DC, that includes classical cDC1 and cDC2. pDC: plasmacytoid DC.

Most of the mature BM cell types overlapped with small populations of thymic cells. The most immature thymocytes, namely DN1 and—to a lesser extent—DN2 cells, overlapped with these mature BM cell types, suggesting a link between immature and mature non-T cells in the thymus (**Fig**. 6D), including B cells, NK cells, pDCs, neutrophils, monocytes, and macrophages (**Fig**. 6C-E and **Suppl. Fig**. S7B-D).

To confirm these findings using a different technique, we used 28-color spectral flow cytometry of human thymocytes to identify mature non-T cell types and their progenitors. Using a principal component analysis to visualize this data in lower dimensions, we again observed a continuum of developing cell types, ranging from immature CD34^+^ progenitors to mature monocytes and B cells, in both the BM and thymus datasets (**Fig**. 6F). Moreover, this data confirms the recently reported development of pDCs in the thymus ^21, 22^. With respect to NK cells, although we identified mature NK cells in both the BM and thymus (**Fig**. 6E, F) we were unable to identify developing NK cells by flow cytometry due to the lack of suitable surface markers. In addition, although mature erythrocytes were lost during the thawing procedure, we found developing erythroblasts in the BM but not in the thymus. Classic DCs from the myeloid lineage (i.e., cDC1 and cDC2) are present in both the BM and thymus (albeit with more cDC1 cells than cDC2 cells in the thymus), but the differentiation steps for these DCs are unclear due to the lack of suitable markers for their corresponding progenitor cells. With respect to neutrophils, their lineage is clearly visible in the BM, but these cells are virtually absent in the thymus samples. Interestingly, however, mature plasma B cells, mast cells, and eosinophils are all present in the thymus (**Suppl. Fig**. S7E).

### Genomic lineage tracing of non-T cells reveals their thymic origin

In the thymus, the first TCR rearrangements occur in the *TCRD* locus, before T cell commitment. The first delta rearrangement occurs between the Dδ2 and Dδ3 segments and is non-functional, followed by the first functional rearrangement between Dδ2 and Jδ1 ^1, 2^ (**Fig**. 7A). Evidence of Dδ2-Jδ1 rearrangements in any given cell indicates that the cell’s differentiation originated in the thymus. Therefore, we sorted B cells, NK cells, monocytes, cDC1 cells, and pDCs from both BM and thymus samples and looked for the presence of these early rearrangement steps (**Suppl. Fig**. S7F); CD34^+^-enriched cord blood cells were included as a negative control, and CD34^+^-enriched thymocytes were included as a positive control ^1, 2^. The presence of the Dδ2-Dδ3 rearrangement—and particularly the Dδ2-Jδ1 rearrangement—in the NK cells, monocytes, cDC1 cells, and pDCs in the thymus samples indicates that these cells can develop in the thymus; in contrast, these same cell types isolated from the BM lack these rearrangements (**Fig**. 7B).

**Fig. 7.**
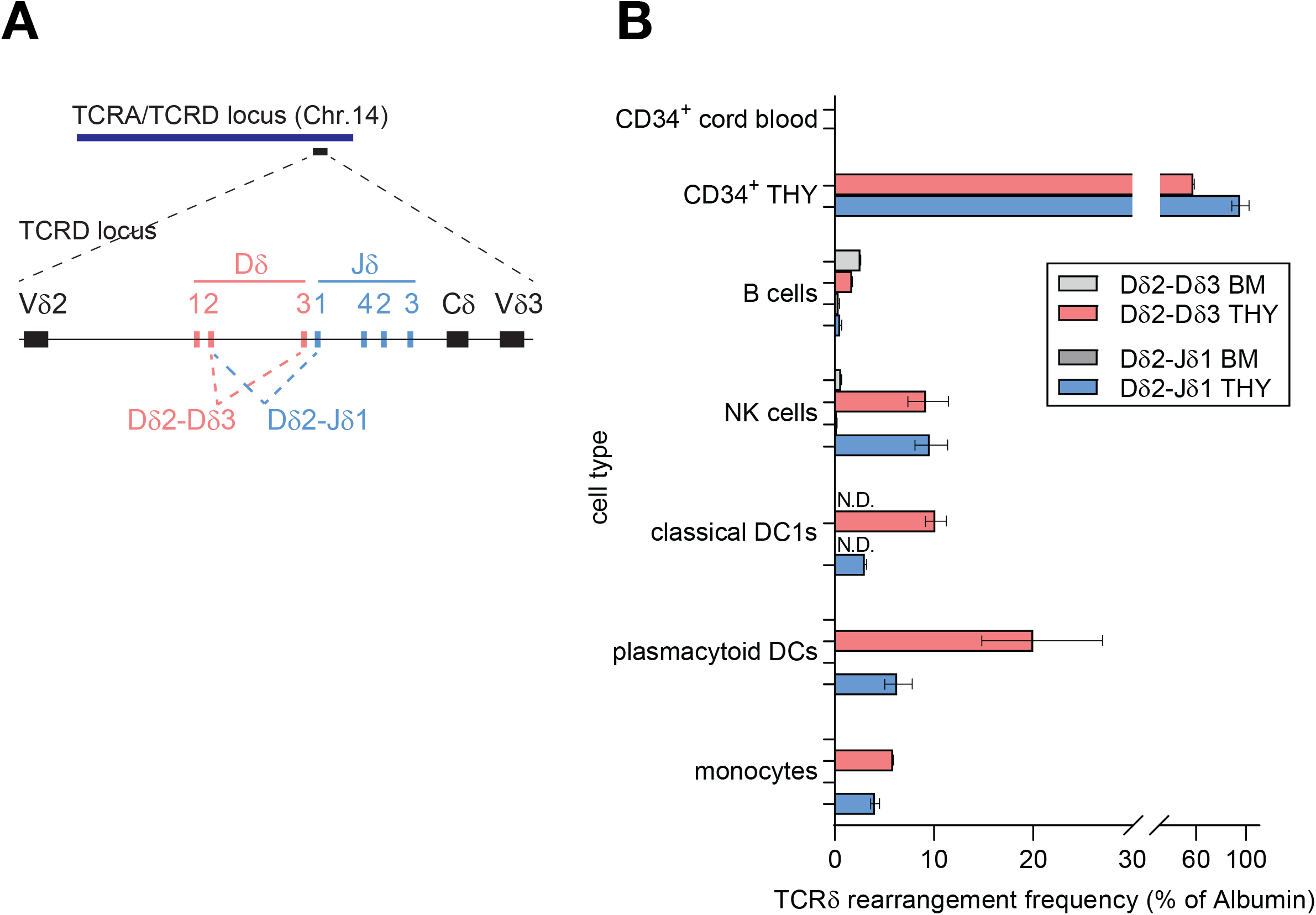
Q-PCR analysis of *TCRD* gene rearrangements non-T cells isolated from thymus and BM. (A) Schematic diagram of the human *TCRD* gene complex. (B) Alleles with Dδ2-Dδ3 (pink) and Dδ2-Jδ1 (blue) rearrangements expressed as a percentage of Albumin in B cells, NK cells, cDC1s, pDCs, and monocytes isolated from thymus, and from BM as controls (light and dark grey, resp.). CD34 cord blood represents a second negative control, whereas immature CD34 thymocytes serve as a positive control.

Interestingly, a small but measurable percentage of B cells in the thymus samples contain the non-functional Dδ2-Dδ3 rearrangement (but not the functional Dδ2-Jδ1 rearrangement); however, the Dδ2-Dδ3 rearrangement is also present at the same frequency in B cells obtained from the BM samples, suggesting that the Dδ2-Dδ3 rearrangement may not be specific to T cells but can also occur in B cells, while Dδ2-Jδ1 rearrangements are extremely rare in B cells. Indeed, a large number of precursor B cell acute lymphoblastic leukemia (precursor B-ALL) cells contain Dδ2-Dδ3 rearrangements, but never contain Dδ2-Jδ1 rearrangements, supporting the notion that the Dδ2-Dδ3 rearrangement marks a step towards the lymphoid lineage that may be common to both T cells and B cells ^54, 55^. Thus, the presence of this rearrangement does not necessarily indicate that the B cells in the thymus developed from thymic progenitor cells. Nevertheless, our combined single-cell transcriptomics, flow cytometry, and *TCRD* rearrangement data provide compelling evidence that non-T cells lineages including—but not necessarily limited to—NK cells, cDC1 cells, pDCs, monocytes, and possibly B cells can develop within the human thymus.

## Discussion

Recent advances in single-cell techniques such as RNA-seq and spectral flow cytometry have led to important new insights into the heterogeneity of the immune system. Here, we used these techniques together with qPCR analysis and functional assays to redefine the road map of T cell development in the human thymus. In our study, we focused on the earliest stages of T cell development, including thymic seeding by progenitor cells and T cell lineage commitment; because we focused on these early stages, these stages are overrepresented in our data; nevertheless, we observed a clear continuum in thymocyte development, with no major gaps in the developmental trajectory.

Using mice with targeted mutations in key transcription factors that regulate T cell development, our group and others previously showed that Tcf7, Gata3, Runx1, and Bcl11b play important roles in this process ^18, 19, 20, 56^. For example, we showed that a transcription factor cascade drives T cell development in which Notch signaling induces its first T cell specific target gene, *Tcf7*, which then induces (in combination with Notch signaling) *Bcl11b* and *Gata3*; *Bcl11b* primarily induces T cell specific genes, while *Gata3* primarily represses non-T cell specific genes ^20, 57^. Interestingly, we observed *TCF7* expression before *BCL11B* expression, suggesting that a similar transcription factor cascade also exists in the human thymus, although carefully designed loss-of-function experiments similar to experiments involving BCL11B ^58^ are needed in order to address this question in the human thymus.

For sorting mature thymocyte subsets, we excluded non-T cell types using lineage markers (except for pDCs, for which sorting markers were not included). Although this strategy allowed us to clearly define the T cell specification and commitment steps, it also made it more difficult to define non-T cell development pathways. However, our CD34^+^-enriched dataset containing DN1, DN2, and DN3 cells did not deplete these alternate lineages and provided important insights into the development of non-T cell types. Moreover, our complementary spectral flow cytometry approach revealed a continuum of cells that develop into several non-T cell lineages in the total thymocyte samples. We also sorted several populations of mature non-T cells from the thymus and confirmed their thymic origin using the earliest *TCRD* locus rearrangements as a lineage-tracing marker. Collectively, our data show that monocytes, NK cells, myeloid DCs, and pDCs can develop in the human thymus. Recently, Lavaert and colleagues reported that pDCs are the only non-T cells that develop in the human thymus ^21, 22^; in addition to confirming this result using lineage tracing, we also observed the development of other non-T cells. We therefore believe that excluding most non-T cells using lineage surface markers during sorting may lead to the false conclusion that only pDCs and T cells develop in the thymus, whereas our detailed scRNA-seq analysis (including immature CD34^+^CD1a^-^ sorted cells) combined with deep flow cytometry profiling and lineage tracing clearly demonstrate that monocytes, NK cells, myeloid DCs, and— albeit to a lesser extent—B cells also develop in the human thymus. Our finding of mast cells in the human thymus is particularly interesting, given that mast cell development in the thymus was reported previously in mice overexpressing GATA3 ^59^, but not in wild-type mice.

Previous studies suggest that an extremely small fraction of thymocytes consists of progenitor cells for NK, B, dendritic, monocytic, and even erythroid cells, in addition to the overwhelming majority of T cell progenitors. Indeed, CD34^+^ thymocyte progenitors can be redirected to differentiate down these non-T cell lineages *in vitro* under the appropriate conditions ^6, 60, 61^ and *in vivo* in humanized NSG mice ^62^. In addition, mouse thymic progenitors were shown recently to develop *in vivo* into B cells and myeloid cells in *Tcf7* knockout mice, which lack T cell development ^20^. Unlike mouse thymocytes, the most immature human thymocytes have an extremely broad differentiation capacity, including the potential to develop into erythrocytes ^6^. However, the fact that progenitors have this capacity does not necessarily mean that this developmental path actually occurs in the human thymus; indeed, despite this inherent capacity we found no indication that erythrocytes develop in the human thymus.

The possibility that B cells can develop in the human thymus has long intrigued immunologists. In mice, clear evidence exists that most B cells in the thymus develop from thymic precursor cells ^63, 64^, although a relatively small percentage of B cells can develop from distinct progenitors, and some mature B cells may seed the thymus ^65^. Using flow cytometry, we previously showed that the full range of B cell developmental stages can be measured in the thymus ^66^; however, this finding that does not exclude the possibility that distinct precursor B cells may have entered the thymus. Indeed, our genetic lineage tracing studies using the initial *TCRD* rearrangements as a marker of T cell progenitors indicate that only an extremely small percentage of thymic B cells are derived from this thymic progenitor. However, it is possible that the TSP3 subpopulation could represent a B/T cell progenitor that follows different developmental trajectories. Nevertheless, the developmental origin of B cells in the human thymus is clearly diverse. With respect to their function, evidence in mice suggests that thymic B cells may play an antigen-presenting role during negative selection ^67^. In addition, evidence suggests that antibody-producing plasma B cells may be present in the human thymus ^68^, a finding supported by our own results.

We characterized the nature of thymus-seeding cells using RNA velocity and RNA-seq data and identified three distinct TSP subpopulations, namely an HSC-like subset (TSP1), an MPP-like subset (TSP2), and a CLP-like subset (TSP3). The HSC-like TSP1 subpopulation appears to be quiescent and may represent a stem cell like reservoir for thymocytes ^69^. On the other hand, the CLP-like TSP3 subpopulation is particularly interesting, as T cells develop much more rapidly from this subset compared to the other two subsets, consistent with previous studies in mice ^70^. Based on marker expression, the TSP3 subset identified here is similar to a previously described CLP subset in human BM ^48^. Recently, Lavaert and colleagues reported two human TSP subsets ^21^, the first of which likely corresponds to our TSP1/TSP2 combined subset due to the expression of *CD34*, stemness genes, and multipotency genes. Our results do not confirm that their second TSP subset is an early thymus-seeding progenitor, as it lacks *CD34* expression but expresses *CD3E*, indicating a T cell lineage committed thymocyte subset. Moreover, their second TSP2 subset likely has robust pDC progenitor potential, as it clustered in our dataset with pDCs and their immediate progenitors; instead, we identified a novel CLP-like TSP subset (which we call TSP3) that develops into committed thymocytes much more rapidly than our HSC-like TSP1 and MPP-like TSP2 subsets.

In addition to characterizing these distinct thymus-seeding populations, we also investigated the T cell lineage commitment for these TSP subsets. We found that compared to the TSP3 subset, the HSC-like TSP1 subset and the MPP-like TSP2 subset have a considerably longer trajectory to becoming committed T cells, characterized by a fully rearranged TCRB (for the αβ lineages) and the expression of *CD3, pTA*, and other T cell specific genes such as *LCK, LAT*, and *MAL*. Interestingly, the branch into the γδ lineage occurs relatively late—from late DN2 to DN3, or even from cells in the ISP or DP stage—in contrast with our previous findings ^1^. In summary, we here report data on the nature of cells seeding the thymus, the potential for development of non T cell lineages, and the establishment of T cell lineage commitment. These findings are important for therapeutic efforts to restore thymic function ^71^, for instance through groundbreaking gene therapy efforts to cure inherited disorders of T cell development, which have recently become clinical reality ^72^ but still need further development to become robust therapies.

## Methods

### Samples

Thymic tissues were obtained as surgical tissue discards from children 7 weeks to 3 years of age (median: 6 months of age) who underwent cardiac surgery; written informed consent was obtained from the parents. The children did not have immunological abnormalities. Thymocytes were isolated from the tissues by cutting the thymic lobes into small pieces and then squeezing the pieces through a metal mesh; the cells were then frozen and stored in liquid nitrogen until further use. Healthy surplus bone marrow mononuclear cell suspensions were obtained from donor bone marrow samples and stored in liquid nitrogen until further use.

### Isolation of thymocyte subsets

To isolate the thymocyte subsets, thymocytes were thawed and pooled from six samples prepared as described above. To isolate DN1, DN2, and DN3 thymocytes, CD34 magnetic beads (CD34 MicroBead Kit UltraPure, Miltenyi Biotec) were used to enrich for CD34^+^ thymocytes.

### Flow cytometry and cell sorting

Spectral flow cytometry and cell sorting were performed at the Leiden University Medical Center Flow Cytometry Core Facility (https://www.lumc.nl/research/facilities/fcf) using a Cytek Aurora 5L (Cytek Biosciences, Fremont, CA) and a BD FACS Aria III 4L (BD Biosciences, San Jose, CA), respectively. To sort the more mature thymocytes, including ISP, DP, and SP cells, non-T lineage cells were excluded using antibodies against the following surface markers: CD26, CD56, CD13, CD33, CD19, and CD34. The purity of the sorted cells is shown in **Suppl. Fig**. S1 and **Suppl. Fig**. S7F, the antibodies used for flow cytometry are listed in **Suppl**. Table S1).

### Single-cell RNA sequencing

Cell suspensions were barcoded (10x Chromium Single Cell platform, 10X Genomics) using the Chromium Single Cell 5’ Library (10x Genomic) from Lin^-^CD34^+^CD38^-^CD1a^-^ (DN1), Lin^-^CD34^+^CD38^+^CD1a^-^ (DN2), Lin^-^ CD34^+^CD38^+^CD1a^+^ (DN3), CD34^-^CD3^low^CD4^+^ (ISP), CD3^-^CD4^+^CD8^-^ (CD4SP), and CD3^+^CD4^-^CD8^+^ (CD8SP) and used to generate eight single-cell 5⍰ gene expression libraries. The loaded cell numbers ranged from 300 to 500,000, aiming for 5000 cells per reaction. All libraries were sequenced using an Illumina HiSeq 4000 (paired-end 75-bp reads) to an average depth of 50,000 reads per cell, resulting in eight datasets (one for each developmental thymocyte subset).

### Paired single-cell TCRαβ/ γδ sequencing

Single-cell TCRαβ sequencing libraries were generated using the Chromium Single Cell 5⍰ Library Construction kit (PN-1000020, 10X Genomics) and the Chromium Single Cell V(D)J Enrichment Kit for human T cells (PN-100005, 10X Genomics). Single-cell TCRγδ sequencing libraries were generated using the Chromium Single Cell 5⍰ Library construction kit (PN-1000020, 10X Genomics) and the following custom primer sets:

1st PCR: forward primer (5⍰ - AATGATACGGCGACCACCGAGATCTACACTCTTTCCCTACACGACGCTC-3⍰) and outer reverse primers for TRDC (5⍰ -GGCAGAAAACCATCAATGCC-3⍰, 5⍰ -GTCTCATTTCTGTTCCTCCC-3⍰, inner reverse primers for TRDC 5⍰ -CCCAGGACTTTTGTCTTCCC-3⍰ and 5⍰ -GAGTGTAGCTTCCTCATGCC-3⍰).

2nd PCR: forward primer (5⍰ - AATGATACGGCGACCACCGAGATCT -3⍰) and outer reverse primers for TRGC (5⍰ - CTCCATTGCAGCAGAAAGCC -3⍰ and 5⍰ - AGACAGCAGGTGATGATGGC -3⍰), inner reverse primers for TRGC (5’- TTCTGGCACCGTTAACCAGC-3’) and (5’-TAGTCTTCATGGTGTTCCCC-3’).

Pooled libraries were sequenced using an Illumina HiSeq4000 (paired-end 150-bp reads), aiming for an average depth of 5000 read pairs per cell.

### Demultiplexing cells to donors

To determine the identity of the individual donor of every cell, we used the Bayesian demultiplexing tool Vireo (v 0.4.2, R version). In brief, we first generated a list of single nucleotide polymorphism (SNP) positions by aligning all expressed reads from all cells and selecting positions with a minimum allele frequency of 0.1 and minimum total coverage of 20. Next, in each cell and at each position we identified overlapping SNPs and counted them in two disjoint groups corresponding to the reference and non-reference alleles. The allelic count matrices were then used to fit a Vireo model that either identified the most likely donor for each cell or classified the cell as a doublet.

### Preprocessing of scRNA-seq data

Sequencing data were demultiplexed and mapped against the GRCh38 genome using Cell Ranger 3.0.0. The resulting gene count matrices were loaded into R using Seurat v3. On average, gene expression was detected for 1000-4000 genes per cell. After investigating the overall gene expression distributions for every sample, we excluded DN1 cells with <100 genes; DN2, DN3, and CD3^-^ DP cells with <200 genes; ISP cells with <150 genes; CD3^+^ DP cells with <85 genes; and CD4^+^ and CD8^+^ SP cells with <75 genes. Low-quality genes were defined as being expressed in fewer than 3 cells. We used the Vireo method based on genotyping to identify the donors *in silico* and used the genotyping profiles to detect doublets. We then corrected the data for low-quality cells and removed cells without a genotype, cells with low gene expression, and doublets. We also corrected for donor effects using canonical correlation analysis (CCA), an integrative method in Seurat version 3.2. Before performing the principal component analysis (PCA), we performed cell cycle regression.

After filtration, we merged the datasets from all 8 thymocyte subsets using the MergeSeurat function. We then split this merged dataset into 6 subsets based on the 6 identified genotypes. We used CCA in Seurat to identify common sources of variation between the donors. After identifying anchors using 40 principal components, the dataset was integrated, and the integrated dataset was used for dimensionality reduction. Before running the PCA dimensionality reduction, cell cycle scores were calculated and used for cell cycle regression based on the expression of genes involved in the cell cycle. The residuals were used to scale the dataset. To maintain the global structure within the dataset as much as possible, we then visualized cells using a two-dimensional UMAP plot generated using the RunUMAP function in Seurat package with 10 principal components. We used the PercentageFeatureSet function to calculate the percentage of mitochondrial gene expression per cell.

Re-analysis was performed on the immature thymocyte populations (“Sort 1”: DN1, DN2, and DN3) and on the more mature thymocyte populations (“Sort 2”: ISP, DP_CD3^-^, DP_CD3^+^, CD4, and CD8). Both datasets started with the raw data and were re-analyzed using the same steps mentioned above. In addition, we removed ribosomal genes, mitochondrial genes, histone genes, and *XIST* (X-inactive specific transcript) from the original count matrix. Low-quality cells were defined as <1000 UMI counts and >30% mitochondrial reads.

After PCA analysis of the Sort 1 dataset, 35 principal components were used for UMAP generation and clustering analysis with the FindNeighbours (dims 1:35) and FindClusters function (resolution = 1), which identified 19 clusters. After scaling the RNA data, the FindAllMarkers function in Seurat was used to identify differentially expressed genes between the 19 clusters. Cluster 19 was identified as a small cluster of stromal cells and therefore excluded from subsequent analysis. Next, FindAllMarkers was again run on the remaining 18 clusters. Re-analysis of the Sort 2 dataset was performed in Seurat using the same parameters as Sort 1, identifying 18 clusters that were used for cluster marker analysis using FindAllMarkers in Seurat.

### Single-cell TCR repertoire analysis

Raw reads from the paired V(D)J sequencing runs were processed using cellranger_vdj in Cell Ranger (v.3.0.0) with a custom reference provided by the manufacturer (version 2.0.0 GRCh38 VDJ-alts-ensembl). V(D)J sequence information was then extracted from the all_contig_annotations.csv output file. Chains that contained the full-length recombinant sequence and were supported by >2 UMI counts were selected and linked to the cellular transcriptome data based on the cell barcodes. These chains were considered productive if a functional open reading frame covering the CDR3 region could be identified.

### Integrated analysis of the thymus and bone marrow datasets

We downloaded the Human Cell Atlas (HCA) bone marrow immune census dataset from https://data.humancellatlas.org/explore/projects/cc95ff89-2e68-4a08-a234-480eca21ce79; this dataset consists of approximately 300,000 cells from eight donors. We removed ribosomal genes, mitochondrial genes, histone genes, and *XIST* from the original count matrix and randomly down-sampled the dataset to 40000 cells (5000 cells per donor). In addition, a CD34-enriched bone marrow dataset was downloaded from (https://data.humancellatlas.org/explore/projects/091cf39b-01bc-42e5-9437-f419a66c8a45); this dataset consists of approximately 30,000 cells from three donors. We also removed ribosomal genes, mitochondrial genes, histone genes, and *XIST* from the original count matrix from this dataset. Both datasets were further processed in Seurat using the same steps used to analyze the thymus data. In addition to the cell annotations provided in the original meta-data, we annotated the cells from both datasets by comparing the differential markers between clusters against cell markers for 35 transcriptionally coherent cell populations ^73^ in the HCA census dataset, and for further sub-classification of cell types we performed differential expression of the annotated cells within the proteo-genomic bone marrow reference maps ^47^.

After cell annotation, we merged both bone marrow datasets with our complete thymus dataset in order to generate a single combined Seurat object. We then performed CCA between our dataset and public datasets using the donor information to identify common sources of variation. We aligned the data using 30 CCA dimensions, regressed out the cell cycle effects, performed PCA analysis, and then generated an integrated 3-dimensional UMAP with 30 principal components. We used BBrowser (version 2.9.23) ^53^ to visualize and analyze the data.

### Automated annotation of TSP subtypes

Version 4 of Seurat introduced weighted nearest neighbor (WNN) analysis as a strategy to integrate multimodal single-cell sequencing data and included a large multimodal BM dataset of 25 cell surface markers with simultaneous whole-transcriptome measurements ^74^. Using WNN analysis to learn the relative utility of each data modality in each cell, this reference dataset contains highly robust cell type annotations. The FindTransferAnchors and MapQuery functions were used to map our DN1/2/3 dataset to the annotated multimodal reference ^74^. In brief, anchors between the query and the reference dataset were identified using a precomputed supervised PCA on the reference dataset to maximally capture the structure of the WNN graph. Next, cell type labels from the reference dataset, as well as imputations of all measured protein markers, were transferred to each cell of the query using the previously identified anchors. Finally, the query dataset was projected onto the UMAP structure of the reference dataset. Using the differential expressed gene markers between the CD34^+^ progenitor populations ^47^, we sub-annotates the CD34^+^ cell types in the BM multimodal reference set ^74^.

### Pseudotime trajectory analysis

For velocity analysis, we generated loom files for each of the 8 datasets using velocyto (version 0.17.17). Using the ReadVelocity function in the SeuratWrappers library, the splicing information from cell cell stored in the loom files was added as new assays to the individual SeuratObjects. We then used the anndata2ri package in Python to convert the Seurat object of Sort 1 (DN1, DN2, DN3) to an AnnData object. This converted all of the results generated in Seurat (including the velocity, PCA, UMAP, and clustering data) to a format that can be used for trajectory analysis with the Scanpy toolkit. Using Scanpy, the count matrix was normalized for each cell and then log-normalized. We used scVelo (https://github.com/theislab/scvelo) to calculate the first- and second-order moments for each cell across its nearest neighbors (scvelo.pp.moments (n_pcs = 30, n_neighbors = 50)). Next, the velocities were estimated and the velocity graph was constructed using the scvelo.tl.velocity function in the dynamic mode and the scvelo.tl.velocity_graph functions. Velocities were visualized using the scvelo.tl.velocity_embedding function on top of the UMAP coordinates that were calculated in Seurat.

For pseudotime ordering of the cells, we used a diffusion pseudotime that infers progression of the cells through a geodesic distance along a diffusion map that was calculated from 10 principal components. From this diffusion map, the diffusion pseudotime (DPT) was computed using the tl.dpt function from Scanpy using 10 diffusion components and setting the root at cluster 1. Together with scVelo we defined 4 developmental paths within the Sort 1 (DN12/3) dataset. We then reordered the cells for each developmental path before generating gene expression heatmaps.

### Analysis of transcriptional regulation

We used SCENIC v. 1.1.2 in R ^40^ to infer the Gene Regulatory Networks in our dataset. SCENIC uses the raw count matrix as the input and filters out genes with fewer than 3 UMIs in 1% of the cells and genes that are detected in at least 1% of the cells. This resulted in a matrix of 220 potential transcriptional regulators from which a gene regulatory network was created using GENIE3/GRNBoost and potential regulons were selected using DNA-motif analysis. We then analyzed the activity of regulons across the cells using AUCell and subsequently visualized the results by plotting the average regulon activity in a heatmap for each cluster in our DN1/2/3 dataset.

### Genomic DNA isolation and qPCR

Genomic DNA was extracted from sorted cell suspensions using the QIAAmp DNA Micro Kit (Qiagen), and DNA concentration was measured using a NanoDrop (ThermoFisher). Quantitative PCR (qPCR) was performed using the TaqMan Universal Master Mix II with UNG (ThermoFisher) in combination with the primers and probes listed in **Suppl**. Table S2 ^1^. All PCR reactions were run on a QuantStudio 3 real-time PCR system (ThermoFisher).

### Cultured artificial thymic organoids

Artificial thymic organoids (ATOs) were initiated with 4000 sorted TSPs or CD34^+^-enriched hematopoietic progenitor cells from cord blood and mixed with 80,000 MS5-hDLL1 cells (Millipore). The ATOs were placed in 0.4-μm cell culture inserts (Millipore) in 6-well plates and cultured in 1 ml/well RPMI-1640 medium (Thermo Fisher) containing 4% B-27 (Life Technologies), 30 μM ascorbic acid (Sigma-Aldrich), 1x penicillin/streptomycin (Gibco), 1x GlutaMAX (ThermoFisher), 5 ng/ml human IL-7 (Miltenyi Biotec), and 5 ng/ml human FLT3L (Miltenyi Biotec) as described previously ^50^.

## Data Availability

All single-cell RNA-sequencing (scRNA-seq) data generated during this study were deposited in GEO: GSE195812 (reviewer token: knmliumadxufziv). This study did not use any unique codes, and all analyses were performed in R and Python using standard protocols from previously published packages. All scripts used in this manuscript will be provided upon request.

## Supplementary Fig. Legends

**Suppl. Fig. S1.**
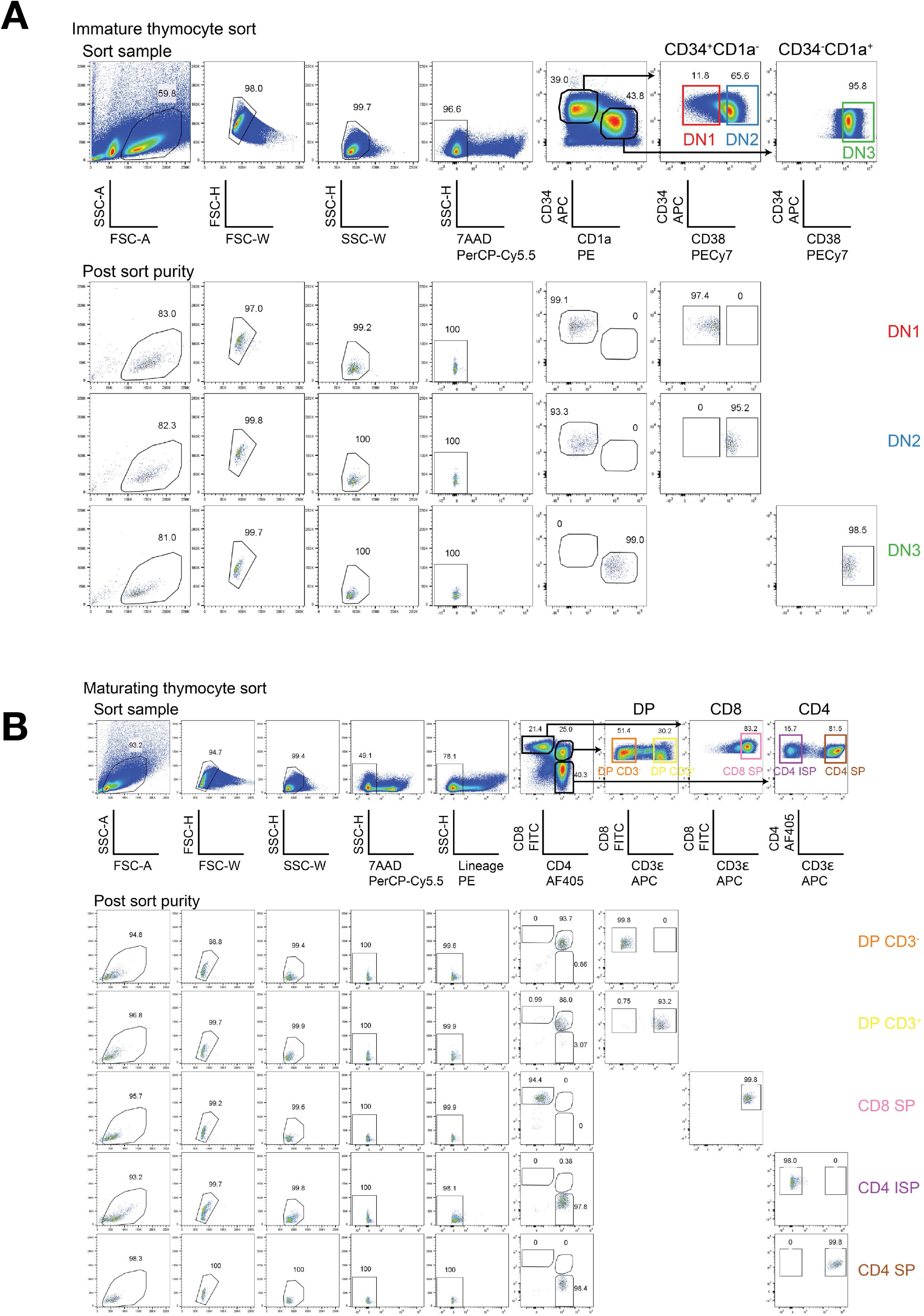
Cell sorting strategies and populations purities. (A) Sorting strategies and population purities of immature DN1, DN2, and DN3 thymocytes. Prior to sorting, thymocytes were enriched for CD34 expression. (B) Sorting and purities of more mature DP CD3 -, DP CD3^+^, CD8 SP, CD4 ISP, and CD4 SP thymocytes.

**Suppl. Fig. S2.**
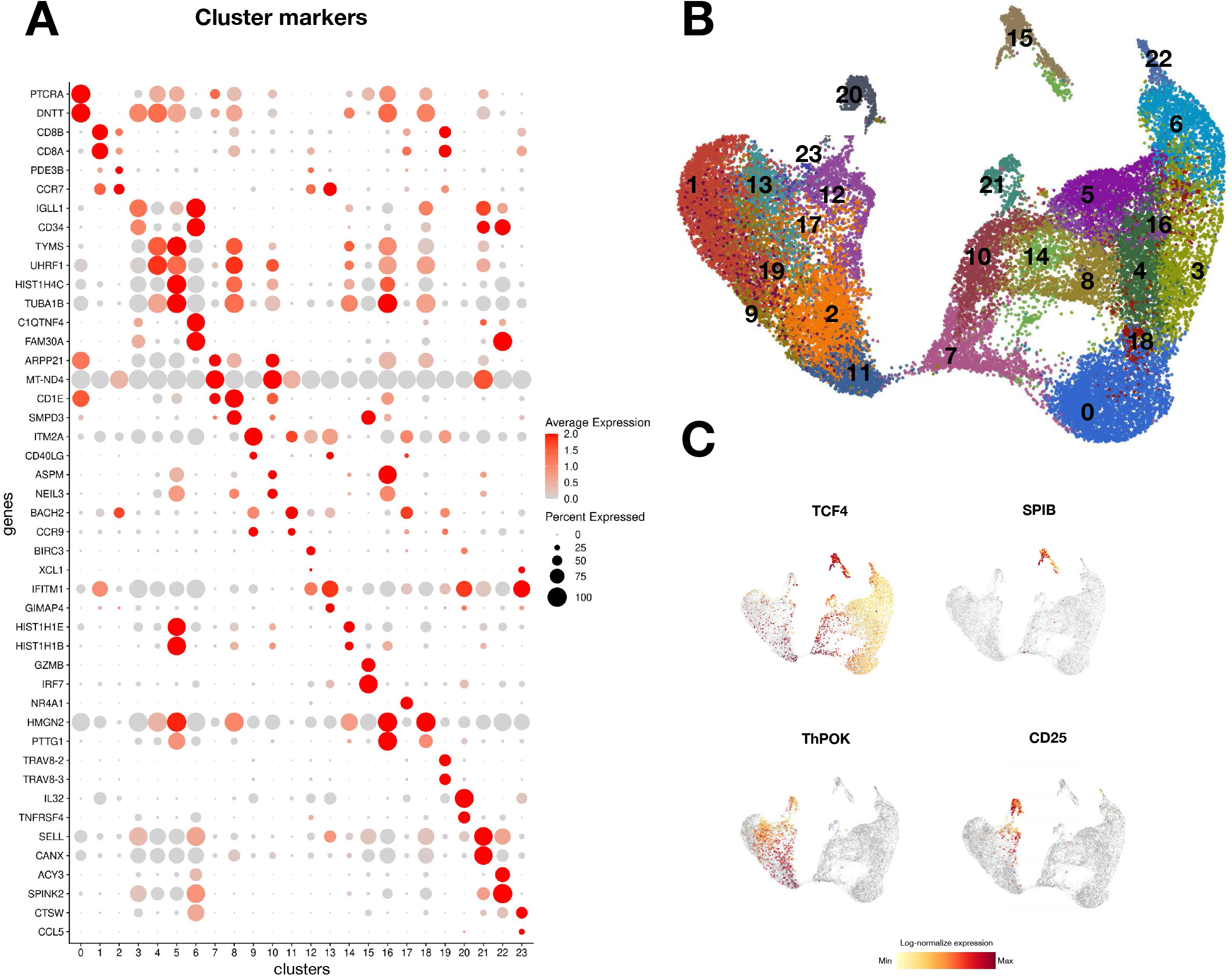
Cluster analysis of the total thymocyte UMAP. (A) UMAP projection of 24 clusters identified by cluster analysis. (B) Dot plot depicting the genes that identify clusters based on differential gene expression. (C) Gene expression of selected genes projected on the total thymocyte UMAP.

**Suppl. Fig. S3.**
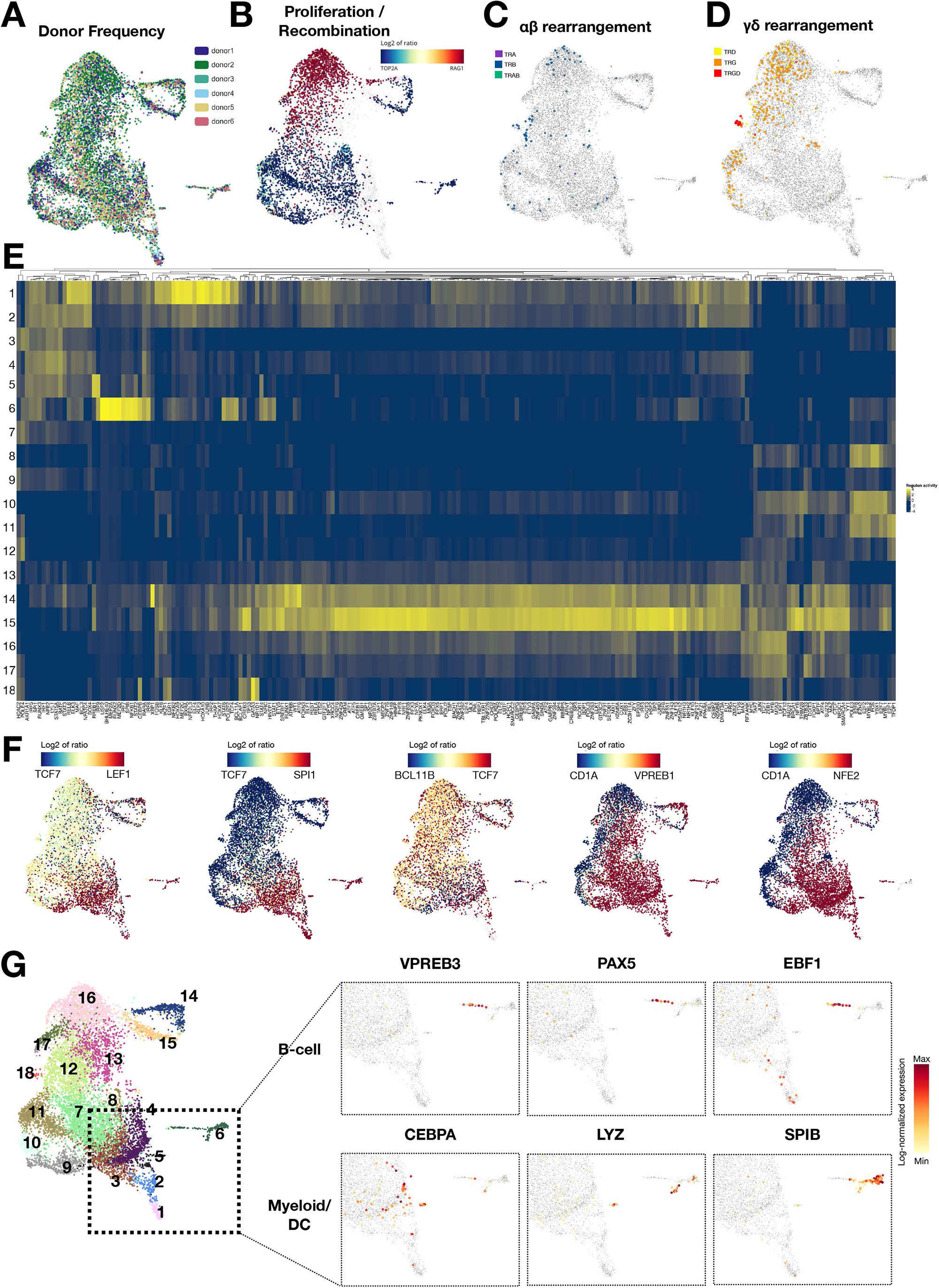
related to Fig. 3. DN1, DN2, DN3 UMAPs with TCR rearrangements and visualization of transcription factor activity and using SCENIC. (A) DN1, DN2, DN3 UMAP visualizing the donor distribution. (B) Proliferation versus recombination activity depicted by expression of TOP2A (blue) versus RAG1 (red). (C) Cells with rearranged TCRα (purple), TCRβ (blue), or fully rearranged TCRαβ (green). (D) Cells with rearranged TCRγ (orange), TCRδ (yellow), or fully rearranged TCRγδ (red). (E) Heatmap of regulon activity as determined by transcription factor activity using SCENIC, within each cluster. (F) Projection of expression of two selected genes (red and blue) per UMAP. Yellow indicates both genes are expressed. (G) Selected gene expression of factors related to B cell and myeloid/pDC differentiation, projected on a partial UMAP with the most immature subclusters.

**Suppl. Fig. S4.**
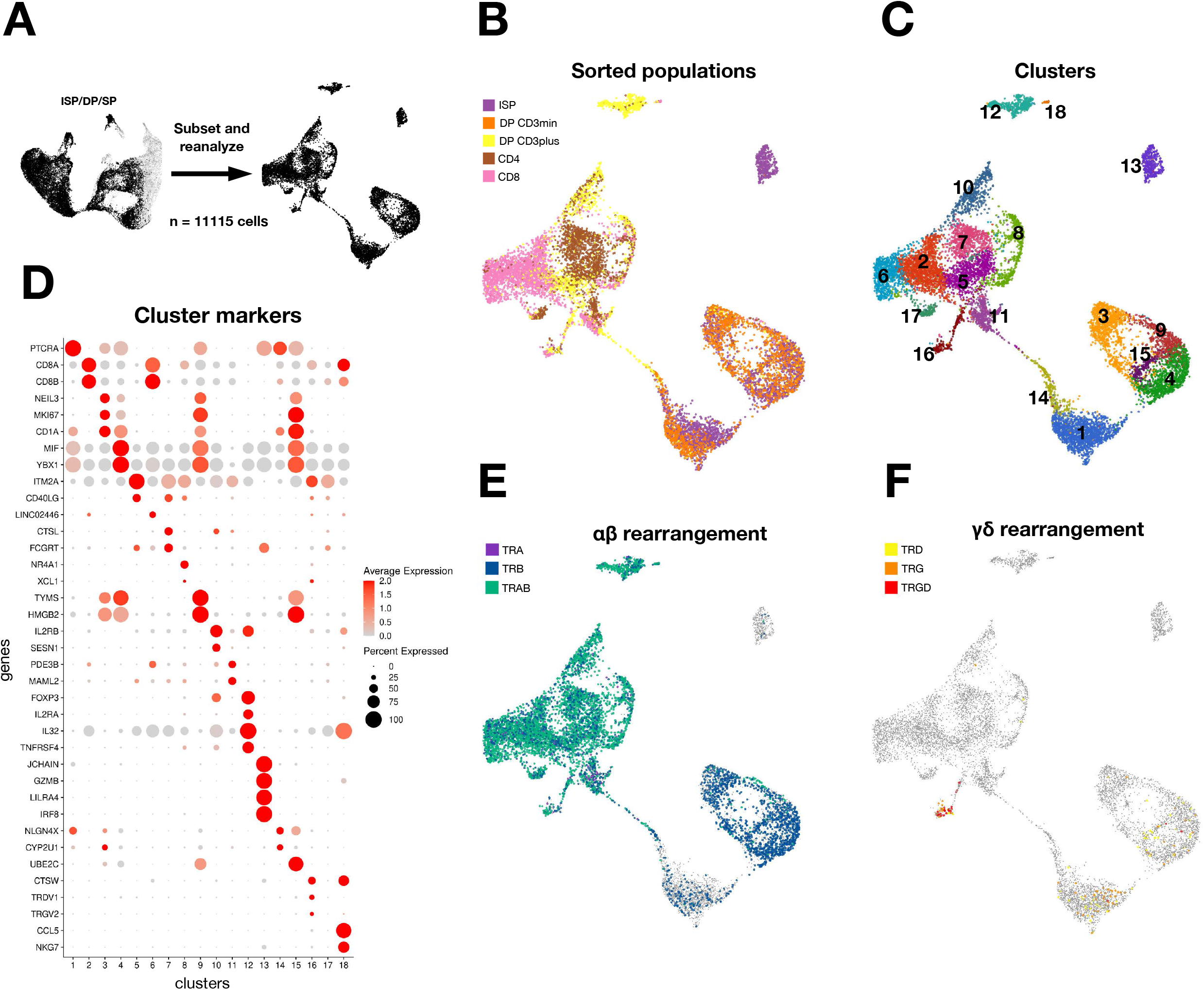
related to Fig. 3. Reanalysis of more mature thymocytes. (A) Reclustering of ISP, DP, and SP cells. (B) UMAP projection of ISP (purple), DP (orange and yellow), CD4 SP (brown), and CD8 SP (pink) cells. (C) UMAP with 18 clusters identified by cluster analysis. (D) Dot plot depicting the genes that identify clusters based on differential gene expression. (E) Cells with rearranged TCRα (purple), TCRβ (blue), or fully rearranged TCRαβ (green). (F) Cells with rearranged TCRγ (orange), TCRδ (yellow), or fully rearranged TCRγδ (red).

**Suppl. Fig. S5.**
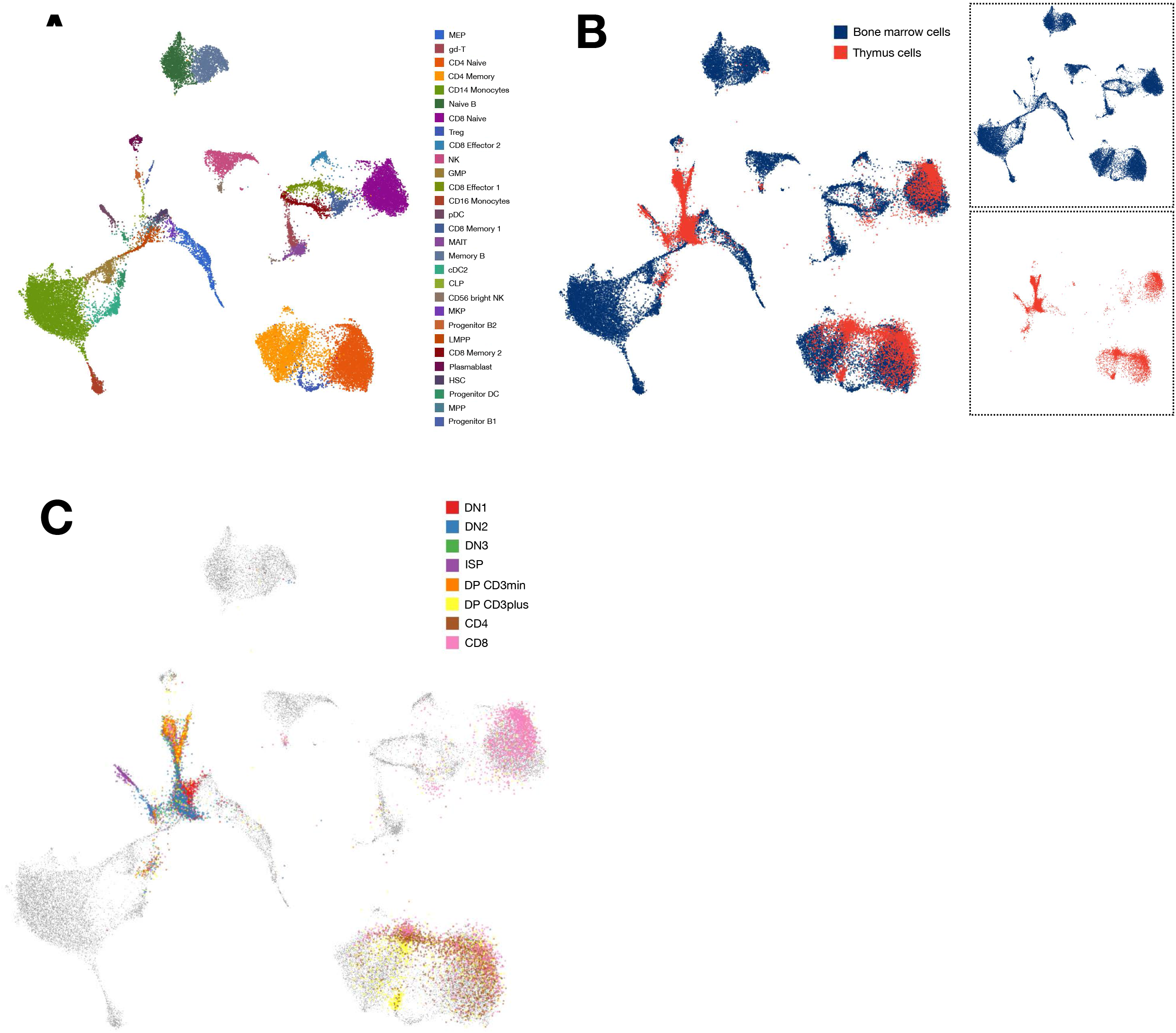
related to Fig. 4. Annotation of thymocytes by multimodal reference mapping. (A) Annotated multimodal BM reference^46^ (B) Total thymocytes (red) mapped onto BM reference (blue). (C) Mapped thymocytes identified by the eight flow-sorted subsets.

**Suppl. Fig. S6.**
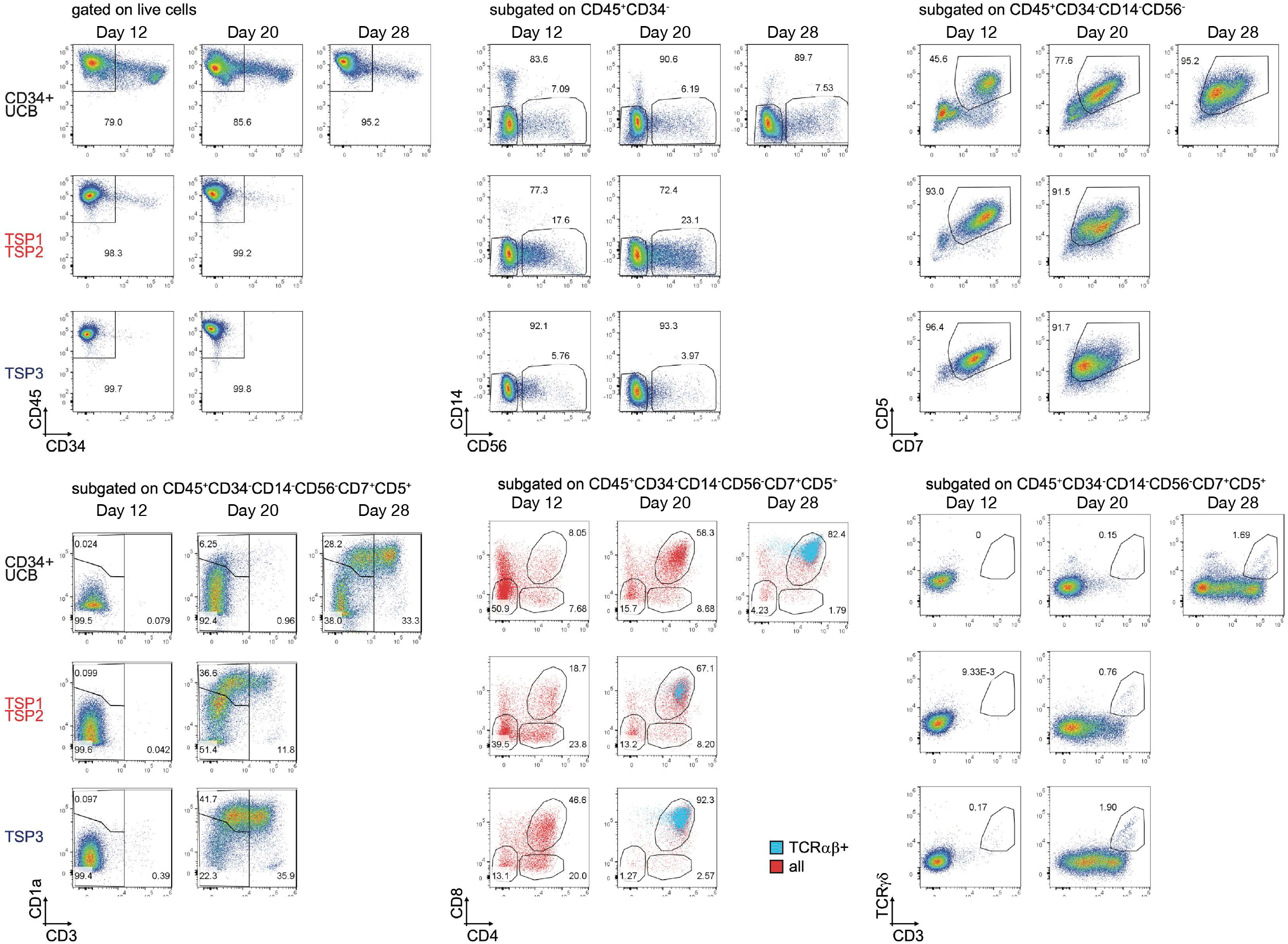
related to Fig. 5D. Additional flow cytometry analysis of TSP cells and CD34^+^ -enriched umbilical cord blood (UCB) cells after 12, 20 and 28 days (only for CD34^+^ UCB) cultured in an artificial thymic organoid (ATO).

**Suppl. Fig. S7.**
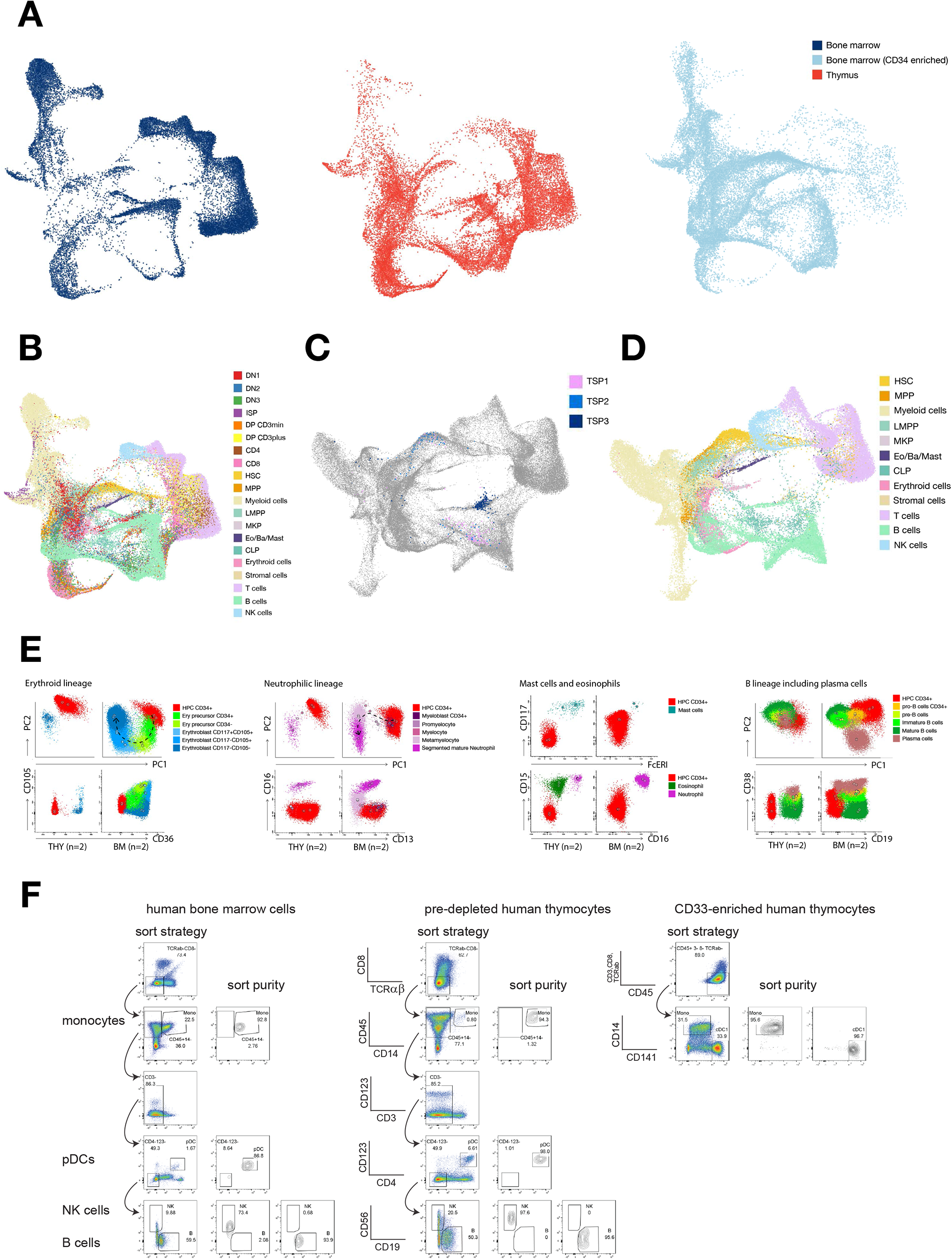
related to Fig. 6. Intrathymic development of alternative lineages. (A) UMAP projection of each individual dataset after integration of mature BM cells ^51^ with CD34^+^ BM ^52^ and our complete thymus dataset (red). with CD34 BM^+ 52^ and (B) UMAP projection of the complete integration of BM and thymus, with annotated BM labels and sorted thymocyte populations. (C) UMAP projection (turned 90 degrees in relation to UMAP in B with Bioturing BBrowser ^53^to demonstrate positioning of TSP1,2,3. (D) BM annotation of cells in the 90-degrees turned UMAP projection. to demonstrate (E) Principal component analyses and example flow plots of human thymocytes using 28-color spectral flow cytometry to indicate mature non-T cell types (erythoid cells, neutrophils cells, mast cells, eosinophils, and plasma B cells) and their progenitors, where applicable. (F) Sorting strategies and population purities of monocytes, NK cells, B cells, pDCs, and cDC1s from human bone marrow and thymus. Prior to sorting, thymocytes were pre-depleted from the majority of T cells using CD8 and TCRαβ (middle panel), or pre-enriched for myeloid cells using CD33 (right panel).

## Acknowledgments

The authors are grateful to the Leiden University Medical Center Flow Cytometry Core Facility for assistance with cell sorting, the Leiden Genome Technology Center, and Janine Melsen for helpful discussions regarding the data analyses.

## Contributions

M.C., K.C.B., F.A.M., C.T., S.M.K, S.V., and L.G.P. performed the experiments. M.C., K.C.B., S.M.K., and C.T performed the analyses. K.C.B., F.A.M., and S.V. provided critical reagents and M.C., K.C.B., E.B.A, K.P.O., M.J.T.R., and F.J.T.S; designed the research and wrote the paper. All authors have seen, reviewed and approved the final version.

